# Nup43 positively regulates *Drosophila* fertility and Myosin VI-dependent actin cone assembly during spermiogenesis

**DOI:** 10.1101/2025.09.29.679220

**Authors:** Jyotsna Kawadkar, Ashley Suraj Hermon, Rohit Kumar, Ram Kumar Mishra

## Abstract

Nuclear pore complexes, composed of nucleoporins (Nups), are critical for bidirectional nucleocytoplasmic transport. Studies on Nups have linked them with various cellular processes, including cell division, contributing significantly to organismal development. Intriguingly, Nup43, an integral member of Nup107 complex, is linked with premature ovarian insufficiency in humans. Here, we report that Nup43 is integral to the maintenance of *Drosophila* fertility. Both the females and males of Nup43 null mutant are sterile. In Nup43 mutants, embryonic development is halted at the first division stage, and males are sterile due to an arrest at the canoe stage of spermiogenesis. The nuclear elongation, shaping, and actin cone assembly steps of individualization complex formation are adversely affected, suspending sperm maturation. Expression through Nup43 transgene in Nup43 null mutants completely rescues spermiogenesis defects. Actin-based motor, Myosin VI (jar), interacts with Nup43 and rescues the actin cone assembly but not the sterility defects. We have uncovered a novel non-canonical function for Nup43 in *Drosophila* spermiogenesis, and propose that Nup43, along with jar, facilitates sperm individualization by promoting actin cone assembly.

## INTRODUCTION

Nucleoporins (Nups) are nuclear envelope-associated proteins that assemble into the nuclear pore complex (NPC). These pores are a dynamic structure responsible for regulating bidirectional macromolecular nucleocytoplasmic transport (NCT) (Petrovic et al., 2022; Beck and Hurt, 2017). Thirty-two distinct nucleoporins, including the ten-membered Nup107-160 complex, organize into subcomplexes and play key roles beyond NCT, like chromatin organization and transcriptional regulation (Lin and Hoelz, 2019; Gozalo et al., 2020; Kuhn et al., 2019; Morchoisne-Bolhy et al., 2015). Nup107 and ELYS/MEL-28, members of the Nup107-160 complex, engage in developmental signaling cascades independent of their traditional functions. In *Drosophila melanogaster*, RNAi-mediated ELYS depletion causes developmental defects attributed to ectopic activation of the Dorsal (NF-κB homolog) pathway even during the late 3rd instar larval stages (Mehta et al., 2020). Nup107 itself has been shown to regulate metamorphosis by modulating the Torso receptor-mediated ecdysone biosynthesis pathway, underscoring its essential role in hormone-driven developmental transitions (Kawadkar et al., 2025). Notably, Nup107 knockdown in somatic gonadal cells induces sterility in females, but males remain fertile (Weinberg-Shukron et al., 2015; Shore et al., 2022). Moreover, Nup107 co-localizes with nuclear Lamin during male meiosis, but its germline depletion disrupts Lamin integrity, resulting in cytokinesis failure (Hayashi et al., 2016).

In *Drosophila*, the early stages of gametogenesis in both sexes have conserved features. The germline stem cells (GSCs) at the anterior tip of the gonad divide asymmetrically to yield one self-renewing stem cell and a differentiating daughter cell. Further, mitotic divisions produce 16-celled structures forming syncytial cysts in both males and females (Bastock and St Johnston, 2008; Demarco et al., 2014). In the ovaries, ovarioles host the egg cell, and its growth and maturation is supported by nurse cells (Bastock and St Johnston, 2008; McLaughlin and Bratu, 2015). In contrast, males produce sperm by a highly coordinated process of spermatogenesis. A 64-cell stage obtained by meiotic division of a 16-cell spermatogonial cyst, round-shaped spermatids are produced. Spermiogenesis is the final phase of spermatogenesis, and involves a series of highly coordinated morphological and biochemical transformations during which round spermatids undergo elongation, cytoplasmic reduction, and nuclear condensation to form mature, motile spermatozoa (Demarco et al., 2014; Fabian and Brill, 2012). During spermiogenesis, actin filament dynamics and branching, ably supported by myosin proteins, play a pivotal role in the individualization complex (IC) formation and motile spermatozoa production. This is an indispensable process, necessary for the separation of interconnected spermatids into distinct spermatozoa (Rogat and Miller, 2002; Zakrzewski et al., 2021).

Beyond organismal development, nucleoporins are being increasingly linked to the development and functional regulation of the reproductive system. Evidence from *Drosophila* and *Xenopus* models highlights the importance of NPC composition and size in oogenesis, influencing NCT during gamete maturation (Feldherr et al., 1998). Mutations in *Drosophila* Nup154, a homolog of mammalian Nup155, affect both females and males. In females, defects range from abnormal nurse cell number, reduced dorsal appendage size, disorganized subcortical F-actin networks, and aberrant apoptotic signaling during oocyte development (Gigliotti et al., 1998; Riparbelli et al., 2007). However, Nup154 mutations in males cause a reduction in testis size, developmental arrest of spermatocytes, and induce complete sterility (Gigliotti et al., 1998). Nup107 complex component Seh1, in coordination with mio, is required for germline cell development, particularly the oocyte maturation (Senger et al., 2011). Nup43 was identified in a clinical exome analysis of human premature ovarian insufficiency (POI), a major cause of female infertility. A loss-of-function mutation in hNup43 is associated with POI individuals (Ke et al., 2023). However, no further analysis was carried out with hNup43, and functional insights into the involvement of Nup43 in reproductive development are limited and necessitate further investigation.

In this study, aiming to investigate the functional role of Nup43 in germline development, we employed CRISPR-Cas9-mediated genome editing to generate a null allele of Nup43 in *Drosophila melanogaster*. Our results demonstrate that Nup43 is indispensable for fertility in both sexes. While the ovarian and oocyte development is unperturbed in Nup43^KO^ females, an arrest of the zygote at the first mitotic division was observed. However, Nup43^KO^ males have disrupted actin cone formation, impaired IC integrity, and individualized spermatozoa production. These defects were fully rescued by germline-specific expression of the Nup43 transgene. While Nup43^KO^ defects have no apparent link with microtubules, the Myosin VI (encoded by jar) expression could restore only the actin cone formation defects. These findings establish Nup43 as an important molecule for gonad development and spermiation events through regulation of actin-cytoskeleton organization.

## RESULTS

### *Nup43* is essential for *Drosophila* fertility

We set out to address the importance of Nup43, an important factor inducing POI-like pathological conditions. We also probed the role of Nup43 in reproductive development and obtained probable mechanistic details in *Drosophila*. We started by comparing the protein sequences of hNup43 with its orthologs in *Drosophila*, mouse, and zebrafish. Our analysis using the protein BLAST (pBLAST) tool revealed that mammalian Nup43 (humans and mice) is highly conserved. They show 93% identity and 97% similarity. However, the conservation is drastically reduced (49% similarity and 26% identity) when humans and *Drosophila* Nup43 are compared (Table S1).

To further explore the function of Nup43 in *Drosophila*, we generated a Nup43 knockout (KO) strain using the CRISPR/Cas9 system. A pair of guide RNAs targeting regions near the start and stop codons of the *nup43* gene was designed to generate null mutants (Fig. 1A). Accordingly, a null mutation was obtained and confirmed through sequencing and PCR analysis (Fig. 1B). The quantitative analysis of Nup43 transcript levels from control and Nup43^KO^ samples revealed a complete lack of ∼ 1500 bp band in Nup43 null mutant, indicating the high efficiency of the Nup43 gRNAs (Fig. 1C). For additional characterization, we produced polyclonal antibodies against full-length Nup43 (refer to the Methods section for details). Immunoblotting of larval brain extracts with custom-generated anti-Nup43 polyclonal antibodies detected a ∼ 43 kDa band in control (Fig. 1D). However, signal for this band was completely lacking in larval brain extracts prepared from Nup43^KO^ animals, confirming the depletion of Nup43 in these null mutants. We immunostained larval salivary gland nuclei with Nup43 antibodies to assess the characteristic nuclear rim staining pattern of Nups. Co-staining with pan-NPC mAb414 antibody (exclusively FG-repeat Nup recognizing antibody) revealed the total absence of Nup43 at the nuclear envelope in Nup43 mutant, without affecting the mAb414 staining pattern (Fig. 1E). Furthermore, we have observed unperturbed nuclear lamina with lamin staining. These observations highlighted the dispensability of Nup43 in lamina and NPC assembly (Fig. S1).

**Fig. 1.**
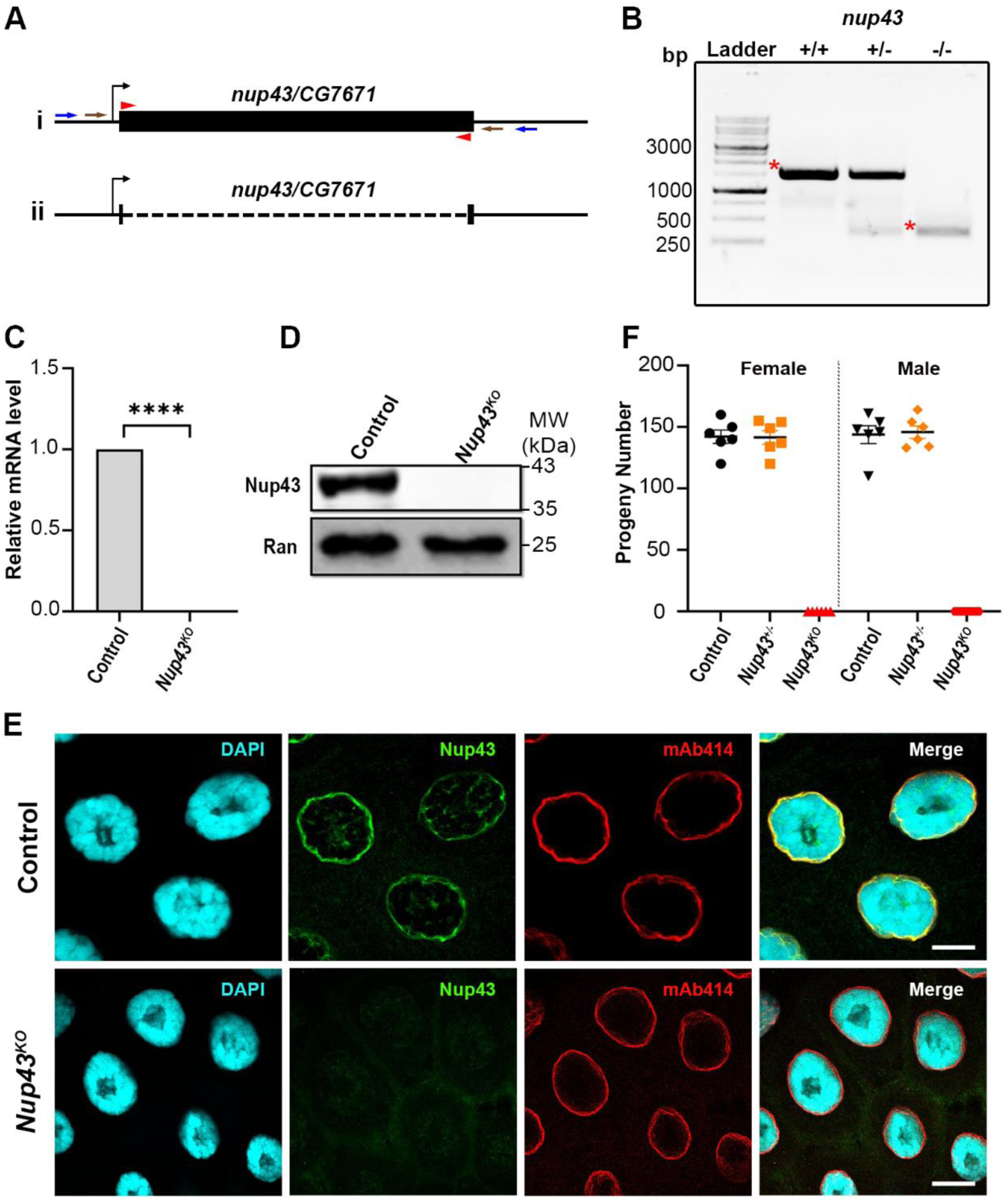
Nup43 is crucial for maintaining fertility. (A) Schematic representation of *Nup43^KO^* generation. (Ai) The genomic locus of *nup43* on chromosome-III (3R). The filled black box corresponds to the *nup43* ORF (1276 bp) with gRNA(s) positions indicated by red arrowheads. The first and second gRNAs were designed near the start and stop codons, respectively, of the *nup43* locus. Blue and brown arrows indicate primer pairs located in the 5’-UTR and 3’-UTR regions used for screening of *nup43* null mutants. (Aii) The black discontinuous line represents the *nup43* (∼ 1210 bp) deletion allele. (B) Confirmation of *nup43* deletion mutant was performed through PCR analysis. In wild-type (+/+) samples, a full-length band (∼ 1500 bp) was observed. In heterozygous (+/-) samples, both the full-length band and the deletion band were present. In homozygous mutant samples, only the deletion band (∼330 bp) was detected. An asterisk (*) denotes the full-length band in wild-type and the deletion band in the Nup43 mutant, respectively. (C) Quantitation of Nup43 transcript levels from control (*w^1118^*) and *Nup43^KO^* larvae. Data are represented from at least three independent experiments. Statistical significance was derived from the Student’s t-test. Error bars represent SEM. ****p = <0.0001. (D) Immunodetection of Nup43 protein levels in third instar larval brain-complex lysates from control (*w^1118^*) and *Nup43^KO^*. (E) Staining of the third instar salivary gland with custom-generated polyclonal anti-Nup43 antibody and Lamin. Chromatin stained with DAPI (cyan) to visualize nuclei. The scale bar represents 10 µm. (F) Fertility analyses of *Nup43^KO^*, males or females of *Nup43^KO^* were mated with *w^1118^* flies, and the number of resulting adult F1 progeny was subsequently counted to assess fertility.

We performed phenotypic characterization, such as morphology and body weight of adults, and rate of adult fly emergence in Nup43^KO^ flies, to observe no apparent defect in flies, and they developed normally as wild-type adults (Fig. S2). It was not a surprising observation as Nup43 has not induced lethality in studies conducted in other organisms. Instead, Nup43 function was implicated in reproductive homeostasis. We asked if Nup43^KO^ adults are fertile. Nup43^KO^ males and females were crossed with w^1118^ flies, and the number of resulting F1 progeny was quantified. Although the Nup43^KO^ adults emerged normally, eggs from Nup43^KO^ did not hatch, and both males and females Nup43^KO^ failed to produce any progeny; thus, they were found to be sterile. In contrast, heterozygous males and females exhibited indistinguishable fertility when compared with control flies (Fig. 1F). These findings imply an essential role for Nup43 in *Drosophila* fertility.

### *Nup43^KO^* embryos fail to complete embryonic mitotic division

To investigate the cause of Nup43-dependent sterility in females, we conducted a series of experiments involving Nup43^KO^ animals. Firstly, we analyzed the morphology of the ovaries of Nup43^KO^ females, and observed no significant changes in the ovariole numbers and their size (Fig. S3). We asked if different aspects of female reproductive development, such as oogenesis, fertilization, and embryonic development, are affected in Nup43^KO^ organisms.

We stained ovaries with Orb, a germline-specific protein highly expressed in post-mitotic ovarian cysts (Lantz et al., 1994), and DAPI to find no defects in oogenesis in Nup43^KO^ females (Fig. 2A). Moreover, when the oocyte development was analyzed in Nup43^KO^ females, normally developing mature oocytes with wild-type-like karyosome were noticed (Fig. S4). This observation is in direct contrast to what has been reported for other nucleoporins, Nup93 and Nup62 (Breuer and Ohkura, 2015). Interestingly, when nurse cells were visualized by Lamin-B staining, their numbers in Nup43^KO^ matched completely with control numbers (Fig. S4).

**Fig. 2.**
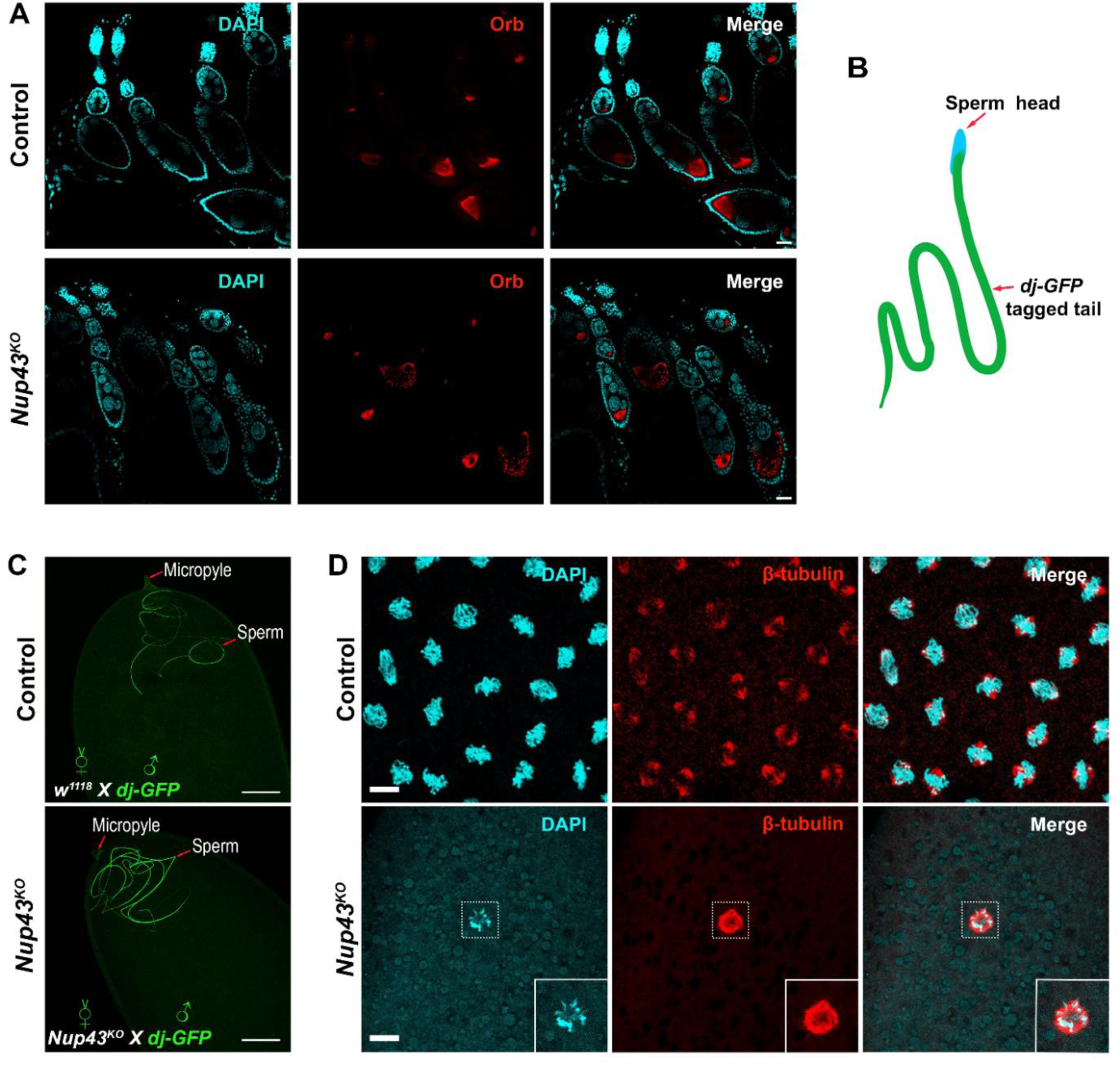
Nup43 is dispensable for oocyte development but essential for the 1st mitotic division after fertilization. (A) Immunostaining of ovarioles from control (*w^1118^*) and *Nup43^KO^* mutants with orb (red) to visualize developing oocytes. (B) Schematic representation of GFP-tagged *Drosophila* sperm. (C) Fertilized egg from control (*w^1118^*) and *Nup43^KO^* mutant females mated with dj-GFP males (labeling sperm with GFP). (D) Immunostaining of the embryos from control (*w^1118^* females mated with *w^1118^* males) and *Nup43^KO^* mutant (*Nup43^KO^* females mated with *w^1118^* males) females, showing microtubules labeled with anti-β-tubulin (red). In relevant panels, Chromatin is stained with DAPI (cyan) to visualize nuclei. The scale bars in (A) is 50 µm, and in (C) and (D) are 20 µm.

Next, we asked if the Nup43^KO^ eggs with normal morphology and micropyle are capable of fertilization. We crossed Nup43^KO^ females with transgenic males expressing sperm Don Juan-GFP (dj-GFP), a flagellum labelling fusion protein (Fig. 2B) (Santel et al., 1997). It is well established that during fertilization, *Drosophila* spermatozoa enter the egg, including the entire very long flagellum (Loppin et al., 2015). Interestingly, the dj-GFP-marked flagellum of the sperm was visible within the egg of Nup43^KO^ (Fig. 2C). This observation confirmed that fertilization occurs in Nup43^KO^ females, akin to wild-type controls.

Thus, the cause of sterility in Nup43^KO^ females should stem from post-fertilization events required for embryonic development. We assessed the progression of fertilized eggs through early mitotic stages of embryonic development in Nup43^KO^ organisms. The early embryonic developmental stages progress rapidly. Approximately 2 h post-fertilization, the eggs from the control had progressed to the syncytial blastoderm stage, with multiple bipolar spindles on metaphase chromatin. However, fertilized eggs from Nup43^KO^ (wild-type males were used for fertilization) exhibited a total arrest at the early chromatin fusion/heterokaryon stages (Fig. 2D). These findings suggest that, while Nup43^KO^ females exhibit normal oogenesis and fertilization processes, they experience a blockage at the first mitotic division during embryogenesis.

### Seminal vesicles of Nup43 mutants are empty and face a sperm drought

We next asked the reason for sterility observed in Nup43^KO^ mutant males. We probed the morphology of the male reproductive system, including testes, seminal vesicles, accessory glands, and the ejaculatory duct. All components of the reproductive system appeared normal in the Nup43^KO^ mutant and were comparable to those of control animals. However, the seminal vesicles (SV) of Nup43 knockout males lacked sperm, were completely empty, and consequently appeared smaller (Fig. 3A,B). This could indicate storage or spermatogenesis defects.

**Fig. 3.**
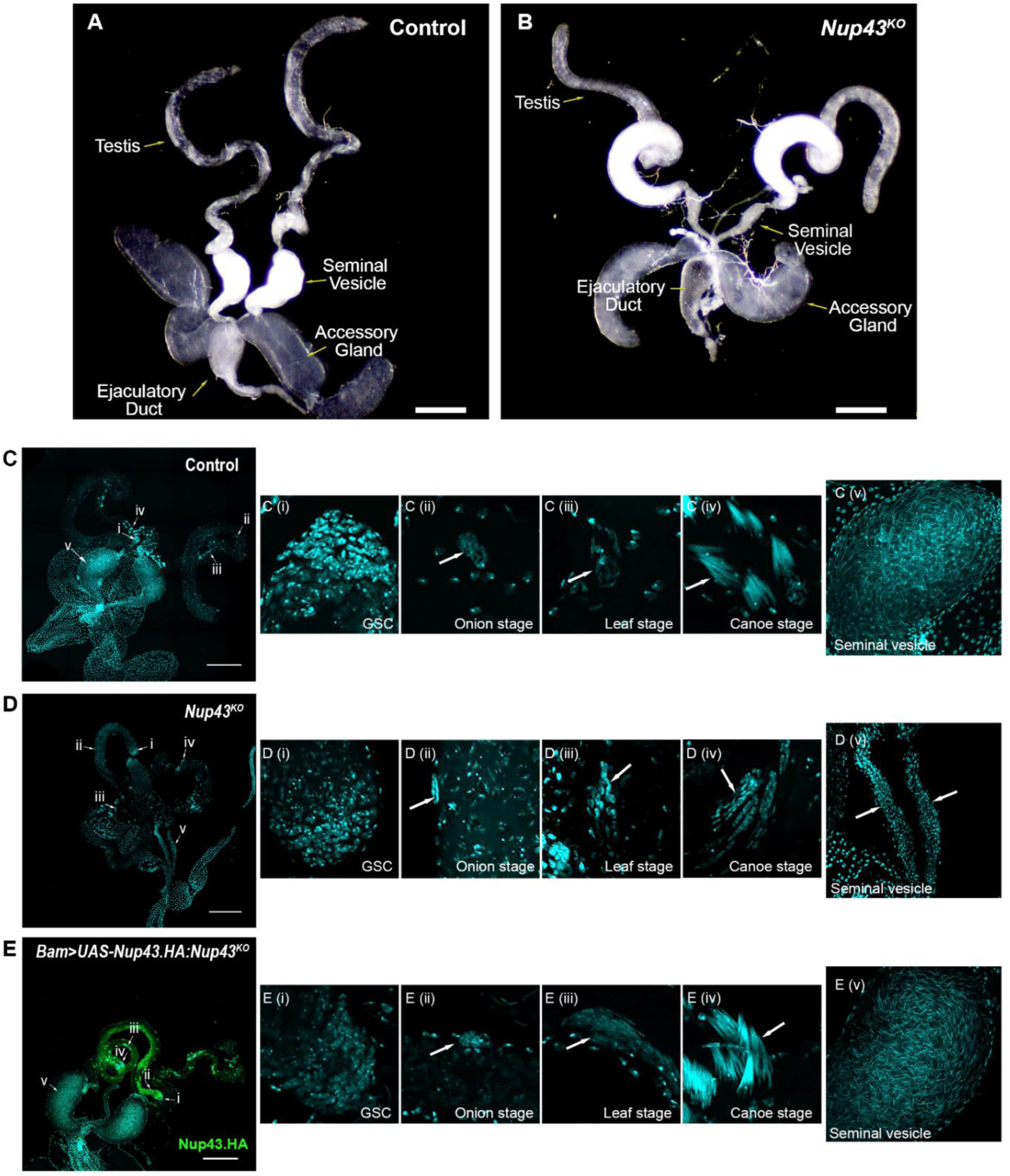
Nup43 is required for the proper progression of spermatogenesis. (A-B) Bright field images of dissected male gonads from control (A, *w^1118^*) and Nup43 knockout (B, *Nup43^KO^*) males. The scale bar represents 300 µm. (C-E) The male testes from control (C), *Nup43^KO^* (D), and rescued animals (E) are shown. Developmental stages progression, germ cell division (i), onion stage (ii), leaf stage (iii), canoe stage (iv), of spermatogenesis are depicted. The stage (v) represents sperm holding seminal vesicles. Fluorescent images (DAPI staining, Cyan) of testes from control males (*w^1118^*), Nup43 mutant males (*Nup43^KO^*), and rescue males (*Bam-GAL4 > UAS-Nup43.HA: Nup43^KO^*). In the (E) panel, expression of HA-tagged Nup43 from a transgene is detected by the HA antibody (green). The scale bar represents 200 µm.

We investigated the spermatogenesis process first, centered around germ cell division. As explained earlier, spermatogenesis in *Drosophila* testes is a multistep process before mature spermatozoa can be stored in the seminal vesicles. In *Drosophila,* sperm development is a highly regulated process, involving communication between germ cells and somatic cells, followed by mitotic and meiotic divisions, differentiation into elongating spermatids, and the formation of testicular bundle structures (Fabian and Brill, 2012; Demarco et al., 2014). We stained testes with DAPI to observe the developmental stages of spermatogenesis. In control animals, all stages of spermatogenesis and spermiogenesis were intact, leading to normal sperm production and its storage in the seminal vesicle (Fig. 3C). In Nup43^KO^ mutants, aspects of spermatogenesis are normal and produce the 64-cell onion structures. However, the Nup43^KO^ mutants are defective in the progression of most of the subsequent steps leading to spermiogenesis. In mutants, spermatids appear to reach only the early canoe stage, where elongation and nuclear shaping of spermatids commence. These spermatids are characterized by aberrations in nuclear shape and elongation. Consequently, spermatids fail to individualize, and no mature sperm is produced (Fig. 3D).

To determine whether the observed phenotype was due to the loss of Nup43 function, we performed a rescue experiment. A transgene coding for 3X HA-tagged Nup43 expression (Nup43.HA) was introduced into the Nup43 deletion background and expressed using the germline-specific *Bam-Gal4* driver. The expression of the 3X HA-tagged Nup43 transgene in germline cells completely rescued the spermatogenesis and spermiation defects (Fig. 3E). Additionally, we confirmed that the rescued males, when crossed with w^1118^ females, produced progenies in numbers comparable to the control as well as heterozygous males (Fig. S5). These findings collectively highlight the essential role of Nup43 in male fertility.

Orb is transcribed in a post-meiotic manner and is translated at the end of the spermatid elongation phase, with high levels found at the caudal end of elongated spermatids undergoing nuclear condensation (Barreau et al., 2008; Lantz et al., 1992; Xu et al., 2012). In control males, the Orb signal is intense and restricted to the caudal end of the testes, representing the tip of the flagellar axoneme. However, in Nup43^KO^ males, the overall Orb signal is diminished and is missing from the caudal end region of the testes (Fig. S6). This suggests a defective organization of the spermatid bundles in Nup43 mutants.

The Don Juan protein is absent from spermatids undergoing elongation, but associates with spermatids in later stages of spermiation (Civetta, 1999; Santel et al., 1997). Accordingly, a dj-GFP-positive mature sperm can be observed inside the seminal vesicles. Lastly, we examined whether the elongation process is impacted in the Nup43^KO^ by assessing the dj-GFP positivity in the seminal vesicles. Dj-GFP positive elongated spermatids comparable to controls were observed in Nup43^KO^ animals. Interestingly, the GFP signal was missing from the seminal vesicles of Nup43^KO^ mutants, in contrast to supersaturating GFP signals in seminal vesicles from control animals (Fig. 4A-B’). The lack of mature sperms in the seminal vesicles was restored to control levels by expression of Nup43.HA in germ cells of Nup43^KO^ males (Fig. 4C-C’). These findings suggest that Nup43 is required for post-meiotic events, including individualization and maturation of sperm during the spermiation process.

**Fig. 4.**
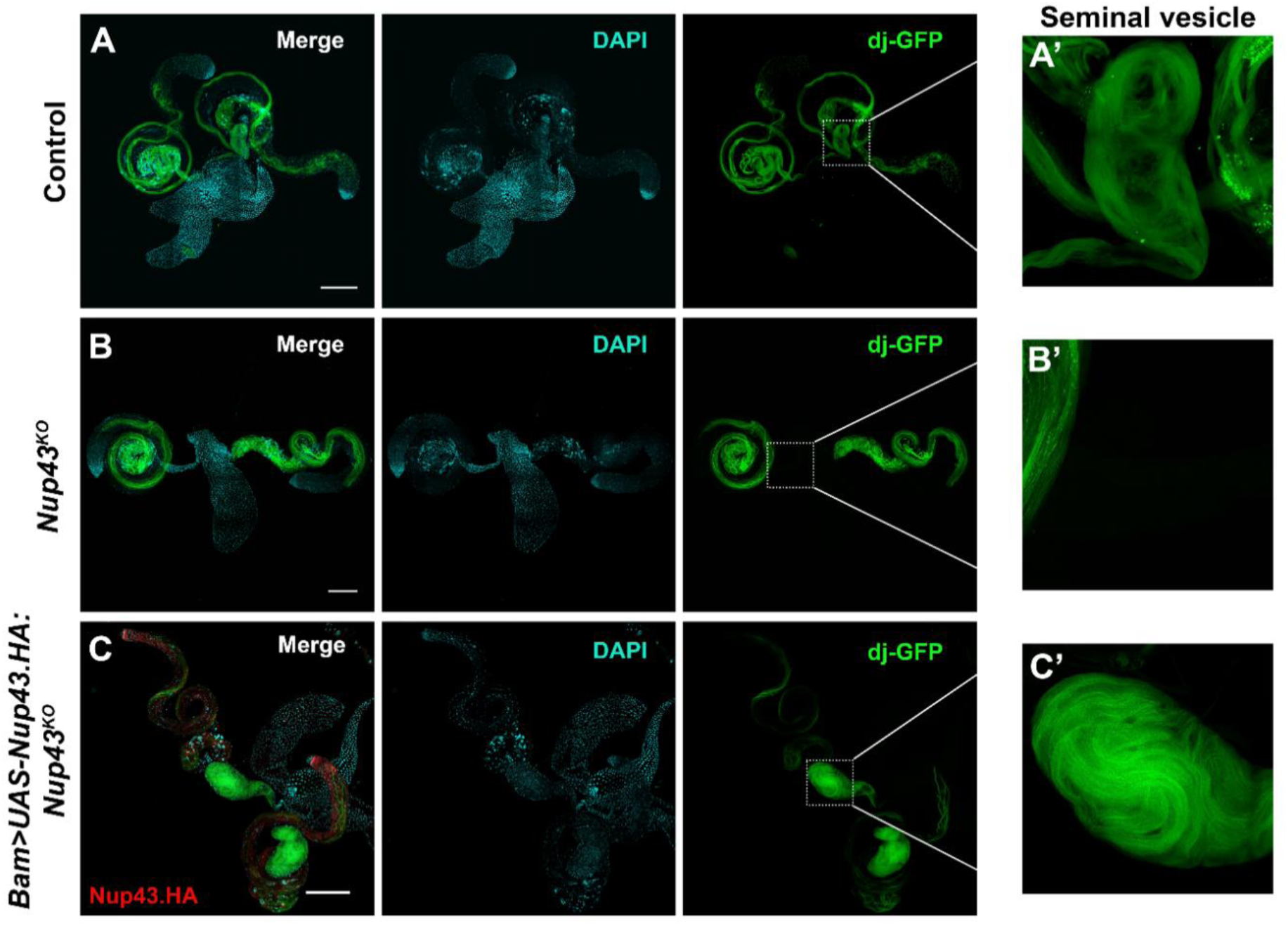
Seminal vesicles in the Nup43 mutant are devoid of sperm. Confocal images showing elongated sperm tails labeled with Don juan-GFP (green) in three groups: (A) control (*dj-GFP*), (B) Nup43 knockout mutants (*dj-GFP; Nup43^KO^*), and (C) rescue males (*dj-GFP; Bam-GAL4>UAS-Nup43.HA: Nup43^KO^*). Zoomed-in views of the seminal vesicle (SV) indicate the presence or absence of GFP-tagged sperm: GFP signal is clearly visible in the SV of control males (A’), absent in *Nup43^KO^* mutants (B’), and restored in rescue males (C’). The expression of the transgene for HA-tagged Nup43 is indicated by the HA antibody, which is shown in red (C, merge panel). Chromatin stained with DAPI. The scale bar represents 200 µm.

### Nup43 is dispensable for nuclear shaping during spermiogenesis

During the final phase of spermiogenesis in *Drosophila* males, the genome undergoes a major transformation for nuclear shaping, and histones are replaced with protamines, leading to a much more compact chromatin structure that’s vital for sperm maturation (Braun, 2001; Oliva, 2006). In *Drosophila*, the male genome is not predominantly associated with histones in a nucleosomal structure; instead, it is organized into a highly compact chromatin structure based on protamines. This unique histone-to-protamine transition occurs during the final stages of spermiogenesis, contributing to chromatin condensation (Braun, 2001; Awe and Renkawitz-Pohl, 2010). In mammals, a similar process begins with the replacement of somatic histones by testis-specific histones during meiosis. These variant histones are subsequently replaced by small, basic transition proteins, which are further substituted by smaller and more basic protamines (Govin et al., 2004; Meistrich et al., 2003; Kimmins and Sassone-Corsi, 2005).

We asked if the histone-to-protamine exchange is impacted in sixty-four spermatids bearing cysts from Nup43^KO^ testes. To check this, ProtamineB-eGFP (ProtB-eGFP) and Histone2Av-mRFP expressing flies were combined with the Nup43^KO^ animals. Interestingly, ProtB-eGFP expressing spermatids and the number of cysts are comparable between Nup43^KO^ and control testes. The ProtB accumulation on chromatin of the developing spermatids is evident, indicating unperturbed Histone-to-Protamine exchange in Nup43^KO^ mutants (Fig. 5A, B). On the contrary, the compact packing and arrangements of spermatid chromatin are perturbed in Nup43^KO^, and they appeared diffused as opposed to a focused pack of 64 spermatids seen in control testes. This observation suggested that Nup43 may be dispensable for early stages of spermatogenesis aspects, but required for their compact arrangement, oriented movement, and spermatozoa formation.

**Fig. 5.**
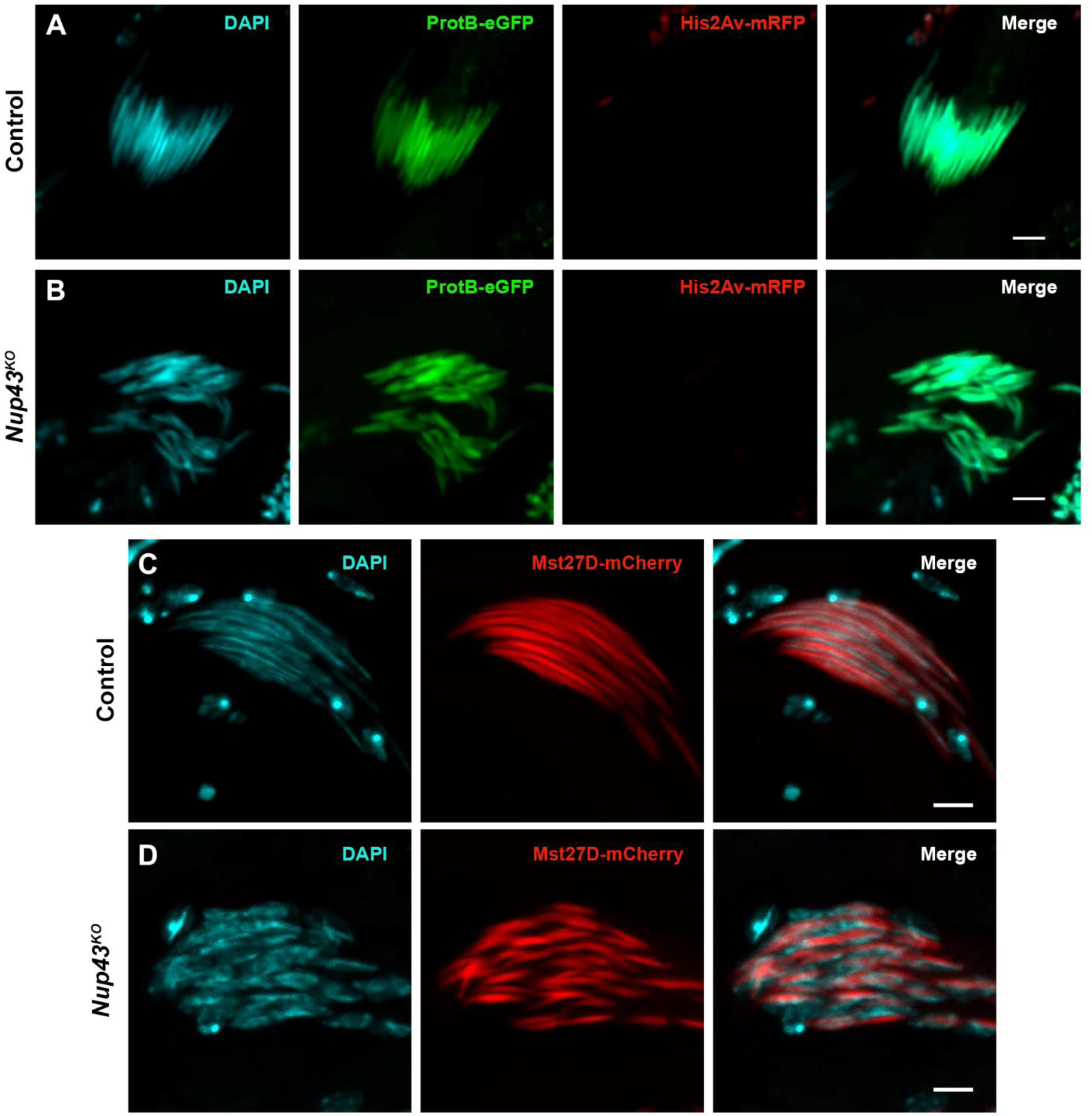
Nup43 regulates normal nuclear elongation during the spermiation process. Fluorescent microscopic images of a spermatid cyst from the testis of control and *Nup43^KO^*. (A-B) Expression and detection of ProtamineB-eGFP (ProtB-eGFP, green) and Histone2Av-mRFP (His2Av-mRFP, red) in the spermatid nuclei of control (A, *ProtB-eGFP: His2Av-mRFP*) and *Nup43^KO^* (B, *ProtB-eGFP: His2Av-mRFP; Nup43^KO^*) testes. (C-D) Spermatids expressing the mCherry-tagged Mst27D (red) in Control (C, *Mst27D-mCherry*) and *Nup43^KO^* (D, *Mst27D-mCherry; Nup43^KO^*) testes. Chromatin is stained with DAPI to visualize the spermatid nuclei. The scale bar represents 5 µm.

During nuclear elongation, the Male-specific transcript 27D (Mst27D) governs the transition of the spermatid nucleus from a spherical to a needle-like shape. Mst27D, through its interaction with the Nup358 component of the NPC, promotes attachment between the nuclear pore complex/nuclear envelope (NPC/NE) and microtubule network to form the critical dense complex (Li et al., 2023; Gartner et al., 2019).

We asked if the perturbations in the nuclear shaping of Nup43^KO^ spermatids can be mediated by Mst27D. We combined mCherry-tagged Mst27D and Nup43^KO^ and examined the testis and spermiation process. Surprisingly, in the Nup43^KO^ background, the Mst27D expression and its asymmetric localization on chromatin are comparable to control levels (Fig. 5C,D). However, the compact structure of the spermatids remains perturbed, and sperm individualization is absent in Nup43^KO^ males, suggesting that Mst27D is not the elusive component of Nup43-dependent male sterility.

Actin-based microfilament and tubulin-based microtubule dynamics play a critical role in the spermiation process in *Drosophila*. The microtubule dynamics are mediated by the efficient functioning of microtubule-binding proteins (Gartner et al., 2019; Nehlig et al., 2017). We asked if the microtubule dynamics is perturbed in Nup43^KO^ mutants. We combined GFP-tagged EB1 (microtubules growing-end tracker) with Nup43^KO^, and investigated the spermiation process in the testes. Similar to the Mst27D observation, the expression and localization of EB1-GFP in the Nup43^KO^ background were unperturbed and comparable to control levels (Fig. S7). Despite this, the individualization of spermatids did not occur, indicating that microtubule networks may not be involved in Nup43-dependent spermiation processes.

### Actin-based Myosin VI motor permits a partial rescue of Nup43 loss-induced spermatid individualization defects

In the final stages of spermatogenesis in *Drosophila*, the elongated spermatid structure is reshaped into individual sperm cells through a process called individualization. At the start of this process, actin cones assemble around the nuclei of the spermatids and then move together in a coordinated fashion from the head region down toward the tail, helping to separate each spermatid into a distinct, mature sperm cell (Fabrizio et al., 1998; Tokuyasu, 1974). Sperm individualization involves membrane remodeling, precise organization of actin structure, and its unidirectional movement over a considerable distance (Noguchi and Miller, 2003).

In wild-type flies, actin cones typically begin to form near the apical side of the spermatid nuclei just before individualization begins. After the nuclei condense, F-actin starts to gather at the junction between the basal and apical regions of the nuclei, with some overlap at the edge of the nuclear envelope (Fig. 6A). As the actin accumulation continues, nuclear condensation completes. To assess if the morphology of F-actin cones and actin organization is affected in Nup43^KO^, we stained control and Nup43^KO^ mutant testes with phalloidin. In a stark contrast to control, actin cones failed to assemble and organize at the front end of canoe-shaped spermatids from Nup43^KO^ males (Fig. 6B). The actin cone-related defects were restored to control levels with expression of Nup43.HA transgene in the Nup43^KO^ background using the *Bam-Gal4* (Fig. 6C). This data suggests that the actin organization (filaments and branched) in maturing spermatids has an important Nup43-dependent component.

**Fig. 6.**
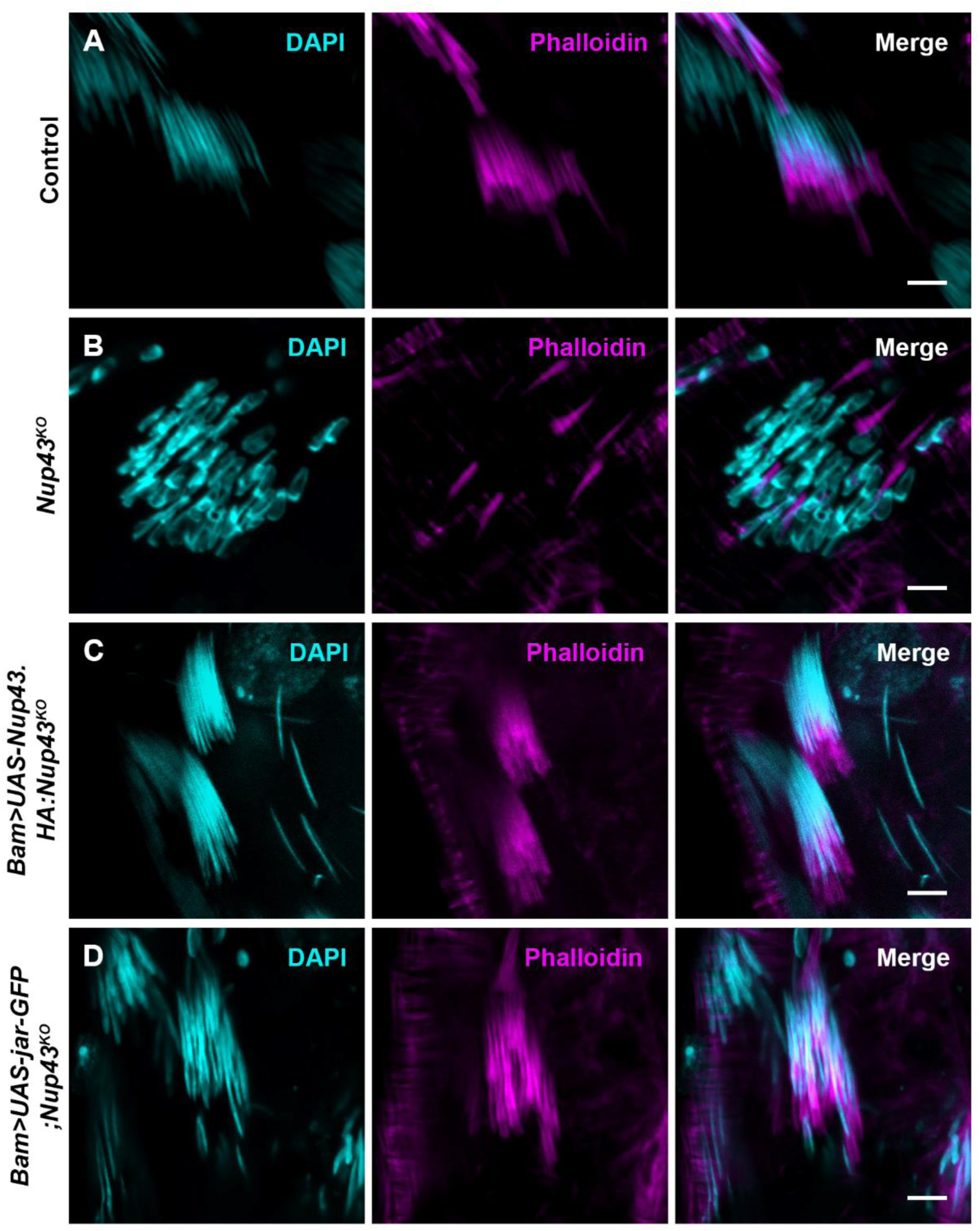
The formation of actin cones is impaired due to the loss of Nup43 during spermiogenesis. Actin cones are marked by phalloidin (magenta). Canoe stage nuclei show the formation of actin cones in control (A, *Bam-Gal4>w^1118^*), Nup43 mutant (B, *Nup43^KO^*), rescue with HA-tagged Nup43 transgene (C, *Bam-Gal4>UAS-Nup43.HA: Nup43^KO^*), and rescue with Jar (D, *Bam-Gal4>UAS-Jar-GFP; Nup43^KO^*). Chromatin is stained with DAPI to visualize the spermatid nuclei. The scale bar represents 5 µm.

After examining the defect in actin cone assembly, the localization of Nup43 during the individualization complex (IC) formation was investigated. During IC complex formation, microtubule organization pulls nucleoporins and nuclear lamina into a dense complex, anchoring organelle-rich cytoplasmic bodies removed as spermatids mature into compact, needle-like sperms. The IC formation is critical as it results in elongated, needle-like sperm production (Li et al., 2023). Surprisingly, Nup43 signals are localized at the base of the 64-spermatid cyst in control testes (Fig. S8). This data provided a hint that Nup43 may interact with some proteins that regulate actin organization during sperm individualization.

During the process of sperm individualization, Myosin VI acts as a dimer to help organize and maintain the cone-shaped actin structure by stabilizing its branched form. In *Drosophila*, the Myosin VI orthologue protein called Jar stabilizes this actin network. When a bunch of 64 spermatids is individualized into a sperm, actin polymerizes, and new branching is generated by the Arp2/3 complex (Noguchi and Miller, 2003; Noguchi et al., 2006). Incidentally, Nup43 was found to bind with Jar during pull-down analysis (Finan et al., 2011), suggesting a potential interaction between the two proteins. To assess the impact of Jar, we combined to express the Jar in germ cells using *Bam-Gal4* in Nup43 mutants. Our analysis showed that when jar was expressed in the recombinant testes (quantified transcripts by RT-PCR; Fig. S9), it allowed actin-cone formation normally (Fig. 6D), but the deposition of sperm in seminal vesicles and rescue of fertility were still missing in jar-GFP expressing males. These results indicate that Nup43 and jar collaborate to help build the actin cone, but do not restore the sperm individualization defect. Nup43 may be assisted by yet to be known molecular players and mechanism(s) for fertility.

## DISCUSSION

Mature sperms are generated following an Actin and microtubule cytoskeleton-based, highly coordinated series of events. One of the key morphological changes in this process is seen with the nucleus changing from round to an elongated, needle-like shape. This process of sperm individualization in *Drosophila* is called spermiation and possesses an additional challenge that needs to be overcome. Nucleoporins (Nups), which constitute nuclear pore complexes, come together in definite stoichiometry and combinations are essential for nuclear shape maintenance. Many Nups have been associated with gametogenesis processes and gonadal dysgenesis defects in different organisms (Gigliotti et al., 1998; Senger et al., 2011; Weinberg-Shukron et al., 2015). Recently, a loss-of-function mutation in human Nup43 (hNup43) was identified in an exome analysis of patients with premature ovarian insufficiency (POI) (Ke et al., 2023). Nup43 is a component of the Nup107-160 subcomplex, which is a key structural unit of the nuclear pore complex (NPC). It is symmetrically distributed on both sides of the nuclear membrane and regulates nucleocytoplasmic transport of biomolecules (Lin and Hoelz, 2019; Loiodice et al., 2004; Cook et al., 2007; D’Angelo and Hetzer, 2008). We have employed advanced genetic tools to probe the involvement of Nup43 in *Drosophila* gametogenesis and gonadal development.

CRISPR-Cas9-mediated Nup43 knockout was generated in *Drosophila melanogaster,* and surprisingly, the animal did not show any obvious developmental/morphological defects. But our analysis indicates that Nup43 is indispensable for fertility in both sexes, as Nup43^KO^ mutant organisms are sterile. The Nup43^KO^ females exhibit normal ovary development, oogenesis, and fertilization processes. However, the zygote is arrested at the first mitotic division post-fertilization, suggesting an essential role for Nup43 in early embryonic cell cycle progression (Fig. 2D). Similar observations were reported with the nucleoporin ELYS; loss of function of *elys* causes female sterility. Mutant females of ELYS exhibit impaired first cleavage division due to abnormal behavior of mitotic centrosomes (Hirai et al., 2018). The halted development may stem from improper structural modifications of the nuclear envelope and nuclear functions observed in Nup43^KO^ mutants in early *Drosophila* embryos (Zierhut and Funabiki, 2015).

Zygotic development is accompanied by spindle formation, progression through cell cycle stages, and rapid division. Mitotic division-dependent stability and centrosome-to-spindle pole localization of Cyclin-B are very relevant in this context (Huang and Raff, 1999; Wakefield et al., 2000). We observed defects in the arrangement and focusing of the mitotic spindle and chromosomal alignment and segregation in Nup43^KO^ mutants (Fig. S10). It may possibly indicate that Nup43 interacts with microtubules or cell cycle regulatory kinases like polo kinase or aurora kinase. Since Nup43^KO^ mutants are arrested at the first mitotic division, similar to another nucleoporin, ELYS mutant, further investigation is needed to determine the significance of Nup43 in *Drosophila* embryonic development.

On the other hand, in Nup43 knockout males, the defects in the spermatogenesis process seem to induce sterility. In the wild-type testes, sperm cells progress through all stages of development, leaf, canoe, and individualization, before being stored in the seminal vesicles (SVs). Interestingly, the SVs in Nup43^KO^ mutants are empty, indicating sperm individualization and maturation defects (Fig. 3B,D). The empty SVs and lack of individualized mature sperm phenotype were seen in pif1A mutants due to the loss of the cystic bulge towards the end of spermiogenesis (Yuan et al., 2019). This phenotype leads to pathological conditions such as azoospermia. Our observations with Nup43^KO^ mutants also present a case of azoospermia, but germ cells appear to divide and progress to the initiation stage, leading to the formation of 64 round-shaped spermatids (Fig. 3D). Use of elongation stage markers such as dj-GFP (Civetta, 1999; Santel et al., 1997) suggested that elongation was occurring (presence of the dj-GFP signal in the testes). However, the dj-GFP signal was absent from the SVs of Nup43^KO^ mutants (Fig. 4B). Further confirming the importance of Nup43 in the spermiogenesis process and sperm individualization, the defects caused by the Nup43^KO^ were fully restored, including sperms in SVs, by expressing the HA-tagged Nup43 transgene (Fig. 4C).

Nuclear shaping includes compaction of chromatin, bringing a drastic change (∼ 200-fold reduction) in sperm nuclei, and many proteins regulate nuclear shaping and assist in the formation of needle-like elongated spermatids (Fabian and Brill, 2012). The key events of nuclear shaping involve exchanging histones with protamines and microtubule motor-based activity for needle-like nuclear architecture (Jayaramaiah Raja and Renkawitz-Pohl, 2005; Awe and Renkawitz-Pohl, 2010; Baarends et al., 1999). While Nup43^KO^ mutants have no mature and individualized sperm, their Protamine exchange (assessed by the use of ProtamineB-GFP line) is unperturbed (Fig. 5B). Mst27D-dependent microtubule organization is important for nuclear elongation, and Mst27D mutants do not lose completely but are impaired in their nuclear elongation (Li et al., 2023). Similarly, in Nup43^KO^, the structure of spermatids remains disorganized, incomplete nuclear elongation, and sperm individualization fails to occur, similar to Mst27D mutants (Fig. 5D). These observations indicate a role for Nup43 in nuclear shaping and elongation steps of the spermiation process, independent of microtubule network and chromatin compaction.

Actin cone (an actin-based branched filamentous structure) at the sperm tail is crucial for the spermiation process and sperm individualization. The actin cone assembly is unperturbed in the Mst27D mutants (Li et al., 2023), and we probed the status of actin cones in Nup43^KO^ mutants. There is no actin cone assembly in Nup43 mutants, thereby impairing the individualization complex (IC) integrity. Through this observation, we emphasize that in *Drosophila* males, Nup43 is essential for spermatid individualization, particularly through the formation of the actin-based IC. Complete rescue of these defects by expression of the Nup43 transgene in Nup43^KO^ mutants suggests a role for Nup43 in nuclear elongation as well as actin cone formation.

Actin cone formation during *Drosophila* spermatogenesis is mainly regulated by the Arp2/3 complex and its key activator, cortactin. This actin assembly is further stabilized by the motor protein Myosin VI, which is encoded by the jaguar (jar) gene (Noguchi et al., 2006; Rogat and Miller, 2002; Noguchi et al., 2008). Notably, a previous study identified Nup43 as a potential interactor of Myosin VI, suggesting a possible functional relationship between these two proteins (Finan et al., 2011). The sperm individualization process relies heavily on proper actin cone formation. Our observations with Myosin VI (jar) in the Nup43 mutant background suggest a practical rescue of actin cone formation defects (Fig. 6D). Nonetheless, sperm individualization and the sterility defects could not be rescued, even partially.

The essence of our findings indicates that Nup43 is critical for the spermiation process, failing which, sperm individualization and maturation aspects are perturbed, leading to sterility (Fig. 7). Interaction of Nup43 with Myosin VI seems to be a critical step in actin cone formation during the spermiation process. Perhaps, additional interactions of Nup43 allow subsequent steps of regular nuclear elongation, nuclear shaping, and sperm individualization. However, the exact mechanism by which Nup43 influences these steps remains unclear and demands further elucidation. Investigation can be directed to identify stage-specific and tissue-specific interactors of Nup43 to shed more light on this. This will enable dissecting the combined role of motor proteins and nucleoporins in regulating cytoskeletal dynamics during gamete formation and maturation. Additionally, this study opens new pathways for understanding the broader roles of nucleoporins beyond their canonical functions, particularly in reproductive development.

**Fig. 7.**
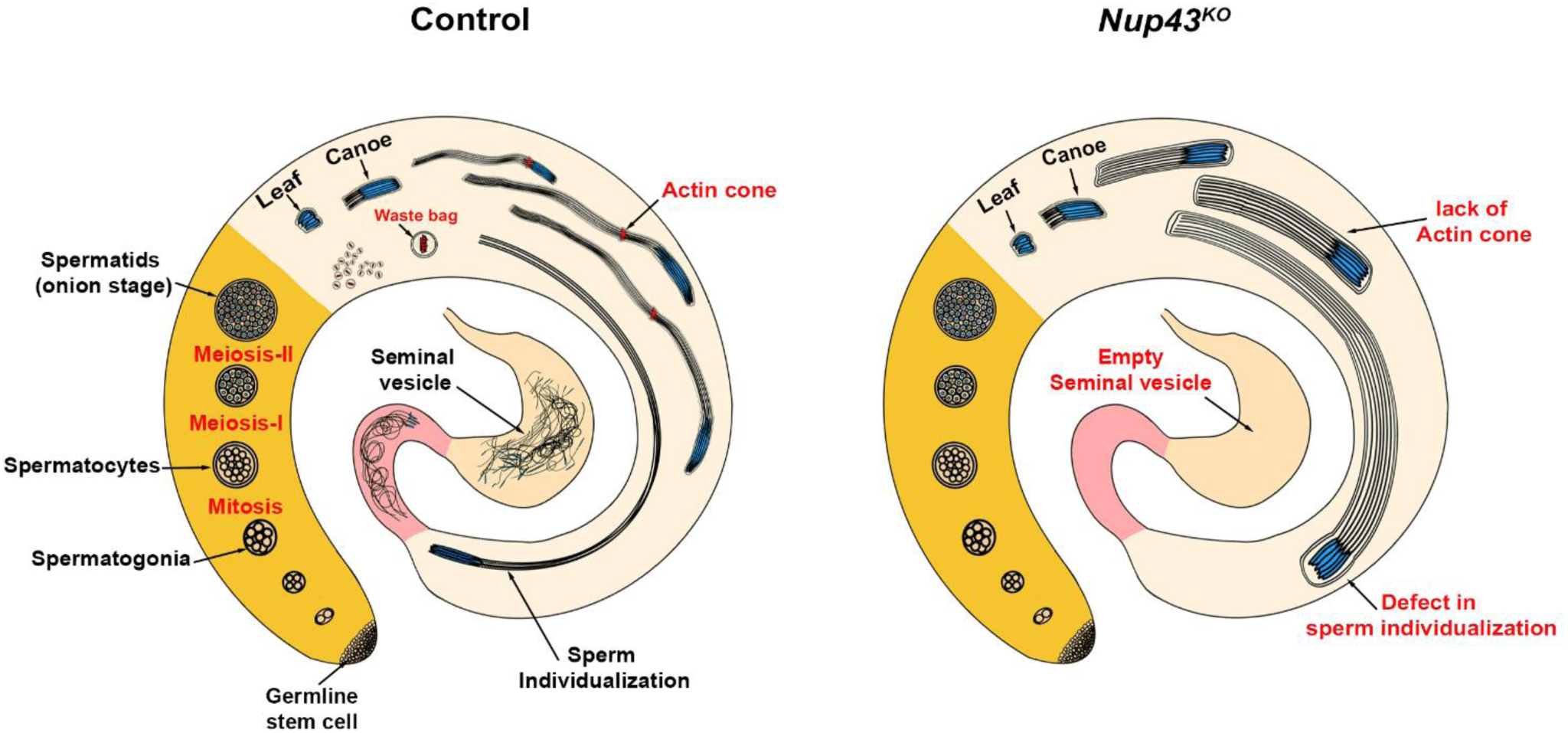
A model for functional regulation of Nup43 on the Spermiation process in *Drosophila* testes. Schematic representation highlighting the spatial progression of spermatogenesis (zone in yellow colour) in the control testis (not to scale). Four mitotic divisions of these germ cells at the apical tip result in interconnected cysts of sixteen spermatogonia. Subsequently, two rounds of meiosis produce 64 haploid round spermatids, referred to as the "onion-stage" of sperm development. Events post-meiotic divisions observe significant morphological changes in developing spermatids. Beginning with the leaf stage and progressing to the canoe stage, then elongating with the help of the actin cone, and ultimately maturing into individual sperm (zone in pink colour). These individualized sperms are stored in the seminal vesicle. In the *Nup43^KO^* mutants, spermatogenesis aspects seem to be unperturbed (zone in yellow colour), but the events related to spermiation fail to take off like the control. Lack of actin cone formation in *Nup43^KO^* mutants perturbs the nuclear shaping and sperm individualization.

## MATERIALS AND METHODS

### In-silico analysis

gfcvgfThe Nup43 protein sequences across different species -humans, fruit flies (*Drosophila*), mice, and zebrafish—were compared using the Protein-BLAST (p-BLAST) tool available through NCBI. The protein sequence for each species was retrieved from the UniProt database (UniProt, 2023). We then used p-BLAST with its default settings to analyze relatedness (similarity and identity) among closely related Nup43 with human Nup43 (hNup43) orthologues.

### Fly stocks and genetics

Experimental *Drosophila melanogaster* stocks were cultivated under the controlled conditions of 25 °C temperature and 60% relative humidity on a standard cornmeal diet (Nutri-Fly Bloomington formulation). Many fly lines used in this study were obtained from the Bloomington *Drosophila* Stock Centre (BDSC) at Indiana University, and Table S2 has the list of all fly lines used in this study. The Nup43 transgenic line (UAS-Nup43.HA) was received as a gift from Prof. Udai Bhan Pandey, USA (Anderson et al., 2021). As a control, *w^1118^* flies were used, and control groups were generated through crosses between the driver line and *w^1118^* flies, as mentioned in the figure legends. All crosses were maintained at 25 °C.

### Transgenic fly generation

We used a systematic approach for the generation of gRNA-containing transgenic lines. We employed an available online tool to design gRNA at http://targetfinder.flycrispr.neuro.Brown.edu with zero off-targets. We utilized an additional tool available at https://www.flyrnai.org/evaluateCrispr/ to ensure and evaluate the predicted efficiency of gRNAs against the *nup43* gene. The most effective sgRNAs, exhibiting elevated specificity without anticipated off-target effects, were chosen and subsequently cloned into the pCFD4 vector. Insertion of the gRNAs in the flyline and generation of transgenic lines were done at the Fly Facility services at the Centre for Cellular and Molecular Platforms, National Center for Biological Sciences (C-CAMP-NCBS), Bangalore, India. Sequences of gRNAs are mentioned in the Table S2.

### CRISPR –Cas9 mediated mutant Generation

gRNA transgenic flies and germline-specific *nanos.Cas9* (BL-54591) were used to create a null mutant of Nup43. The first cross was done between virgins of *nanos.Cas9* flies and males of gRNA transgenic lines, and males of the F1 generation were collected. In the second cross, F1 males were crossed with balancer flies of the third chromosome. F2 progeny were utilized for the single-line cross (with balancer flies) to assess the deletion of *nup43*. Subsequently, F3 progeny were subjected to nested PCR to confirm deletions. Positive individuals were reared for the experimental use.

### Genomic DNA Isolation

To isolate genomic DNA for the screening of mutants, a conventional method was used. 250 µL of solution A (0.1 M Tris-HCl, pH 9.0, 0.1 M EDTA, and 1% SDS, supplemented with Proteinase-K) was used for homogenization of ∼ ten adult flies, followed by incubation at 70 °C for 30 min. Subsequently, 35 µL of solution B (8 M potassium acetate) was added, mixed gently without vortexing, and incubated on ice for 30 min. To get the clear supernatant, the homogenate was centrifuged at 13,000 rpm at 4 °C for 15 minutes, and the supernatant was cautiously collected without disturbing the pellet. 150 µL of isopropanol was added to the supernatant and incubated on ice for 15 minutes. After centrifugation at 13,000 rpm for 5 minutes, the supernatant was discarded, and the pellet was washed with 1 mL of 70% ethanol by centrifuging at 13,000 rpm for 5 minutes. Again, the supernatant was discarded, and the pellet was dried at 55 °C until the ethanol evaporated. Finally, 50 µL of Tris-EDTA buffer was used to dissolve the dried pellet and incubated at 37 °C.

### Nested PCR

To assess the presence of deletions in the mutant fly genome, we used the Nested PCR. We started with 1 µL of genomic DNA as the template for the initial PCR, which utilized an outer set of primers (shown in Fig. 1A, marked in blue arrow in the UTR region). After this, 1 µL of the initial PCR product was used for a second PCR with an inner set of primers (shown in Fig. 1A, marked in brown arrow in the UTR region). The resultant PCR products were then separated on a 0.8% agarose gel and analyzed for DNA bands representing the deletion (∼330 bp) mutation. The primers used in this study are listed in Table S2.

### Generation of recombinant flies

To generate recombinant flies, we have checked the distance between two genes to determine the likelihood of recombination. A greater distance between genes increases the chances of crossing over. We performed a cross between one fly containing the Nup43 mutation and another fly carrying 3X HA-tagged UAS-Nup43 on the third chromosome. After this cross, the F1 progeny were test-crossed to a balancer, resulting in the F2 progeny. These F2 progeny were then used for single-line crosses to assess recombination. Screening for recombinants was conducted using PCR-based methods. The same method was used for the generation of His2Av-mRFP and ProtB-eGFP recombination on the second chromosome.

### Antibody generation and western blotting

To produce antibodies against *Drosophila* Nup43 (CG7671), the full-length gene was sub-cloned into the modified tag-less pET28a(+) vector, specifically the pET28a(+)-JK vector. The protein expression, purification, and antibody generation protocol were followed as described in (Kawadkar et al., 2025).

For immunoblotting, third instar larval brain complexes were dissected and lysed in 1X Laemmli buffer. Cleared supernatant samples (equivalent to two head complexes) were loaded into each well and separated using 8% SDS-PAGE. Following transfer to a polyvinylidene difluoride membrane, blocking was conducted using 5% fat-free milk. The membrane was subsequently treated with polyclonal anti-Nup43 antibody (1:250) and anti-α-Ran (BD, TC4, 1:5000) at 4 °C overnight. After three washes with TBS-T buffer (20 mM Tris-HCl, pH 7.5, 150 mM NaCl, 0.1% Tween-20), the membrane was incubated with secondary antibodies: anti-rabbit-Alexa-Fluor Plus 680 (Thermo Fisher Scientific, #A32734) and anti-mouse-Alexa-Fluor Plus 800 (Thermo Fisher Scientific, #A32730) for 1 hour at room temperature. Following incubation with secondary antibodies, the membrane was washed three times with Tris-buffered saline containing Tween-20 (TBS-T, 10 minutes each) to eliminate non-specifically bound secondary antibodies. Later, the membrane was used with the Li-Cor IR system (Model: 9120) for imaging, facilitating the observation and characterization of the protein bands on the membrane.

### Embryo processing for immunostaining

Flies were allowed to lay eggs for 20 to 30 minutes on a mixed fruit juice agar plate. Sticking food residues were removed from the embryos by washing them immediately with water. Dechorionation was performed using a 1:1 diluted 4% stock bleach solution until the dorsal appendages disappeared. Later, the bleach solution was removed, and the embryos were washed thoroughly to remove the bleach. Next, fixation of the embryos was performed in 1:1 (formaldehyde: n-heptane) solution for a brief period of 2 to 5 minutes. The formaldehyde layer at the bottom was then discarded, and the embryos were transferred to a 1:1 solution of methanol (100%) and n-heptane, where they were kept for 2 to 5 minutes. The embryos in this solution were aggressively shaken for two minutes in order to remove the vitelline membrane. The middle and upper layers were carefully removed, followed by the methanol layer, leaving just enough to keep the embryos wet. The embryos were dehydrated by washing them in 100% methanol twice. The samples were subsequently preserved at −20 °C in 100% methanol unless needed. For immunostaining, the embryos were rehydrated, and the same protocol was followed as for *Drosophila* tissues (salivary gland and testis).

### Immunostaining of *Drosophila* tissues

The previously reported protocol for immunostaining *Drosophila* third instar larval salivary glands and other tissues (such as testis and ovaries) was followed (Mehta et al., 2021). Dissection of late third instar larvae was conducted in cold phosphate-buffered saline (1X PBS) to isolate the salivary glands. The salivary glands were pre-extracted for 5 minutes in 1% PBST (1X PBS + 1% Triton X-100), followed by fixation in freshly prepared 4% paraformaldehyde for 30 minutes at room temperature. Thereafter, the glands were thoroughly rinsed with 0.3% PBST (0.3% Triton X-100 in 1X PBS). Blocking was conducted for 1 hour using 5% normal goat serum (The Jackson Laboratory, USA, #005-000-001). The glands were incubated overnight at 4 °C with the following primary antibodies: anti-Nup43 (1:100), mAb414 (Biolegend, 1:500), and anti-lamin Dm0 (DSHB, ADL67.10, 1:800). Post-primary antibody incubation, tissues were washed thrice with 0.3% PBST (PBS + 0.3% Triton X-100), and incubated with secondary antibodies: anti-rabbit Alexa Fluor 488 (Thermo Fisher Scientific, #A11034, 1:800) and anti-mouse Alexa Fluor 568 (Thermo Fisher Scientific, #A11004, 1:800) for 1 hour at room temperature. Following the incubation with secondary antibodies, tissues were subjected to three washes with 0.3% PBST (PBS + 0.2% Triton X-100) and were then mounted using DAPI-containing Fluoroshield (Sigma, #F6057). The same protocol was followed for staining the gonads, ovaries, and testes. The gonads were stained with the following primary antibodies: anti-orb (DSHB, 6H4, 1:50), *β*-tubulin antibody (DSHB, E7, 1:400), mouse anti-HA (CST, 2367T, 1:400), and rabbit anti-HA (CST, 3724T, 1:400). Afterward, secondary antibodies were applied, including anti-mouse Alexa Fluor 488 (Thermo Fisher Scientific, #A11029, 1:800), anti-rabbit Alexa Fluor 568 (Thermo Fisher Scientific, #A11036, 1:800), and anti-rabbit Alexa Fluor 647 (Jackson ImmunoResearch, 1:800). Additionally, Phalloidin-647 (Jackson ImmunoResearch, PHDN1-A, 1:400) was used to stain filamentous actin in cones.

### Bright-field microscopy

To capture images of the testis and ovaries of adult flies, gonads were dissected and fixed with 4% PFA and were mounted on slides and imaged using a Nikon Stereo Microscope (SMZ745T) upright light microscope at 10X magnification. For each genotype, at least five individual gonads were analyzed, with samples collected from three separate, independently conducted crosses.

### Fertility Assay

To assess female/male fertility, individual flies were paired with three control flies of the opposite sex (*w^1118^*) and kept at 25 °C. Mating was allowed to proceed for either 3-4 days, depending on the experiment. Afterward, flies were removed, and the vials were incubated further to allow progeny to develop. Following the emergence of the F1 progeny, adult flies were collected, and the average progeny count per vial was documented. The values obtained were plotted in GraphPad Prism and subsequently evaluated statistically using Student’s t-test. For each genotype, ten separate vials were set up, and the entire fertility assay was repeated three times to ensure consistency.

### Fertilization test

Virgin females of the *Nup43^K^*^O^ mutant, aged three days, were crossed with age-matched dj-GFP males (BL-5417). After allowing the flies to mate for 24 hours, eggs were collected during a 30–60 minute window. These freshly laid eggs were immediately processed; they were first rinsed with water, then dechorionated with 1:1 diluted 4% stock bleach solution until the dorsal appendages disappeared. After thorough rinsing with distilled H_2_O, the eggs were fixed for 15– 20 minutes in 4% PFA. After fixation, they were washed with 0.3% PBST and then with 1X PBS. The processed eggs were mounted on coverslips using Fluoroshield mounting medium (Sigma, #F6182) and imaged under a 40X oil immersion objective using the Olympus FV3000 confocal microscope. Z-stack images were captured and stacked to visualize the presence of dj-GFP in the sperm tail, which was used as an indicator of successful sperm entry into the egg.

### Weight analysis

Groups of ten adult flies were weighed using an electronic balance. Three replicates were performed for each strain, control (*w^1118^*) and *Nup43^K^*^O^. Statistical comparisons were made using a two-sided Student’s t-test.

## Acknowledgments

The fly lines used in this study were generously provided by the Bloomington *Drosophila* Stock Centre (BDSC), and we sincerely thank them for their support. We especially thank Prof. Udai Bhan Pandey for providing the fly line for Nup43 overexpression. Special thanks to Pradyumna for creating the *dj-GFP/CyO; Nup43^KO^/TM6.Tb* combination. Special recognition is extended to the Developmental Studies Hybridoma Bank (DSHB) for providing antibodies and to Bio Bharati Life Sciences Pvt. Ltd., Kolkata, India, for their essential contribution to antibody generation. We want to express our gratitude to C-CAMP Bengaluru for their invaluable support in developing the transgenic fly. We acknowledge the Central Instrumentation Facility of the Indian Institute of Science Education and Research Bhopal for invaluable assistance received in DNA sequencing and access to confocal microscopes.

## Author Contributions

JK and RKM conceptualized the project and the experimental plan and wrote the manuscript. JK performed all experiments and analyzed data. ASH and RK helped to perform experiments on oogenesis and the rescue part, respectively. RKM provided intellectual support in data analysis and secured funding for the project.

## Data and resource availability

All relevant data and details of resources can be found within the article and its supplementary information.

