## Supplemental file for "Nup43 positively regulates *Drosophila* fertility and Myosin VI-dependent actin cone assembly during spermiogenesis"

**Supporting Information**

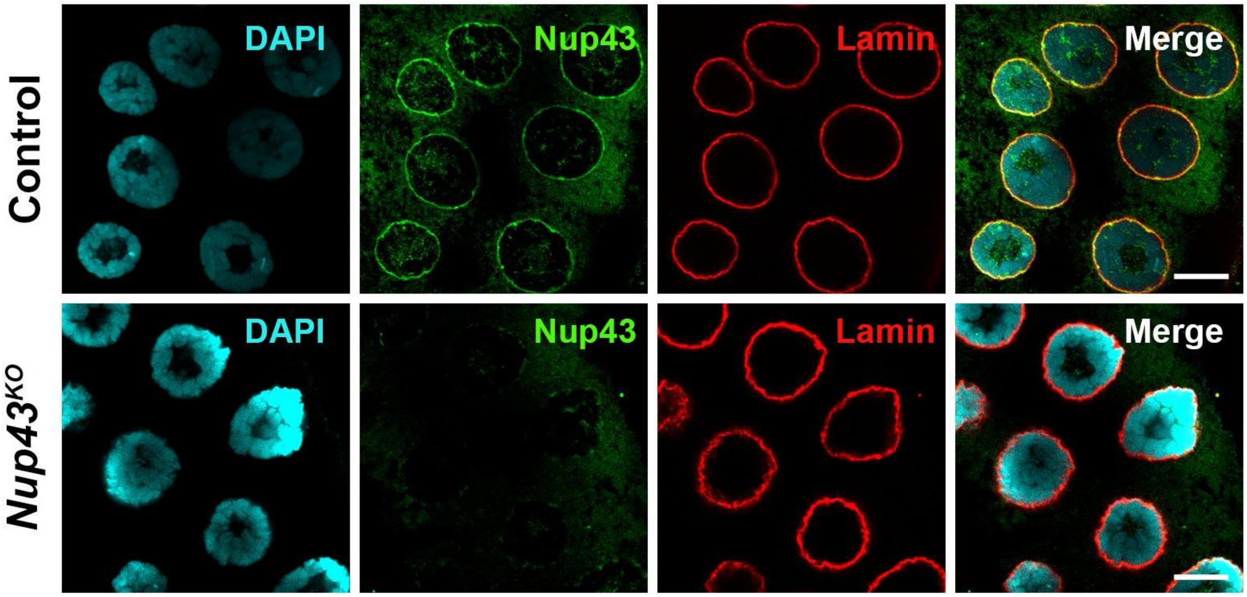

**Fig. S1. Nuclear lamina is unperturbed in *Nup43*$^{KO}$ mutants.** Immunostaining of salivary gland nucleus from control (upper panels) and *Nup43*$^{KO}$ (lower panels). The Nup43 is stained in green, and lamin-B is stained in red. Chromatin stained with DAPI (cyan) to visualize the nucleus. The scale bar represents 10 µm.

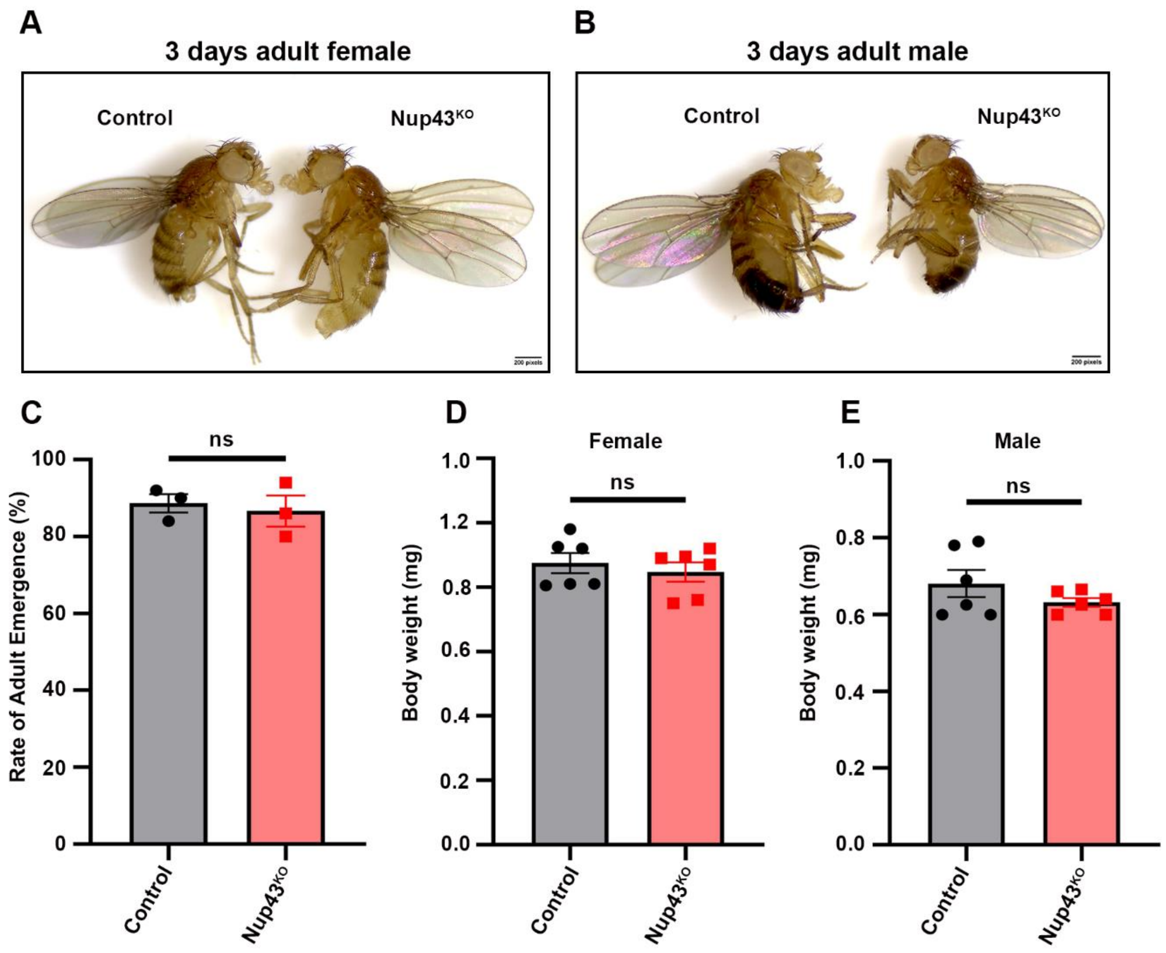

**Fig. S2. Regular morphological features and physiological parameters are unaffected in *Nup43$^{KO}$* mutants.**

(A) Images of a 3-days-old female of the control and *Nup43$^{KO}$*. (B) Images of a 3-days-old male of the control and *Nup43$^{KO}$*. Ten adult flies of each genotype were analyzed (n=3). (C) Rate of emerging adult progeny for control and *Nup43$^{KO}$* adults. A total of ten flies were used for each assay, with three assays conducted (n=3). (D and E) The weight of control and *Nup43$^{KO}$* females (D) and males (E) were analyzed using ten adult flies per genotype (n = 3).

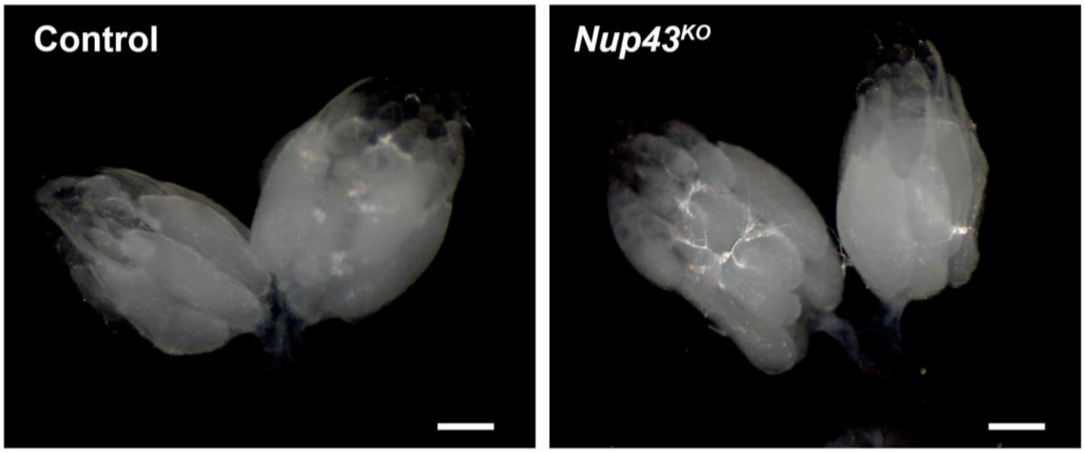

**Fig. S3. The morphology of *Nup43$^{KO}$* ovaries is normal.** Brightfield microscopic images showing the ovaries of 3-day-old adult females with *Nup43$^{KO}$* (right) and control (*w$^{1118}$*, left). The scale bar represents 300 µm.

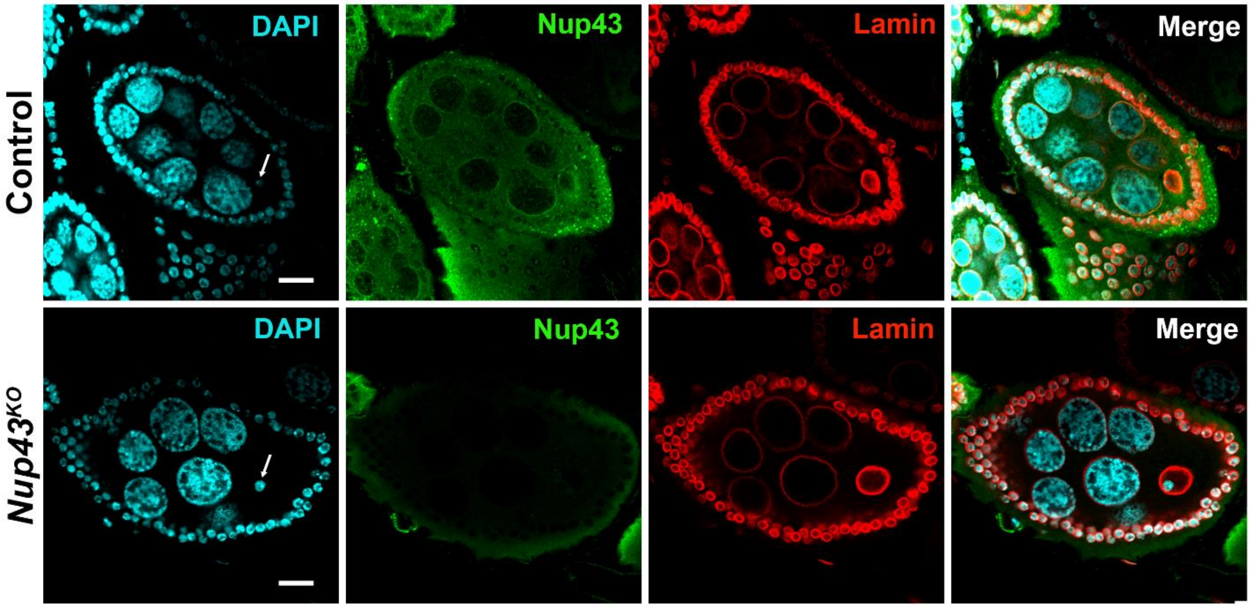

**Fig. S4. Nup43 colocalizes with lamin-B at the nuclear rim of nurse cells and developing oocytes.**

Immunostaining of ovarioles of control and *Nup43*$^{KO}$ with anti-Nup43 (stains nuclear pores, green) and anti-Lamin-B (stains nuclear rim, red) antibodies. White arrows indicate the presence of karyosomes in the oocyte nucleus. Chromatin stained with DAPI (cyan) to visualize the nucleus. The scale bar represents 20 µm.

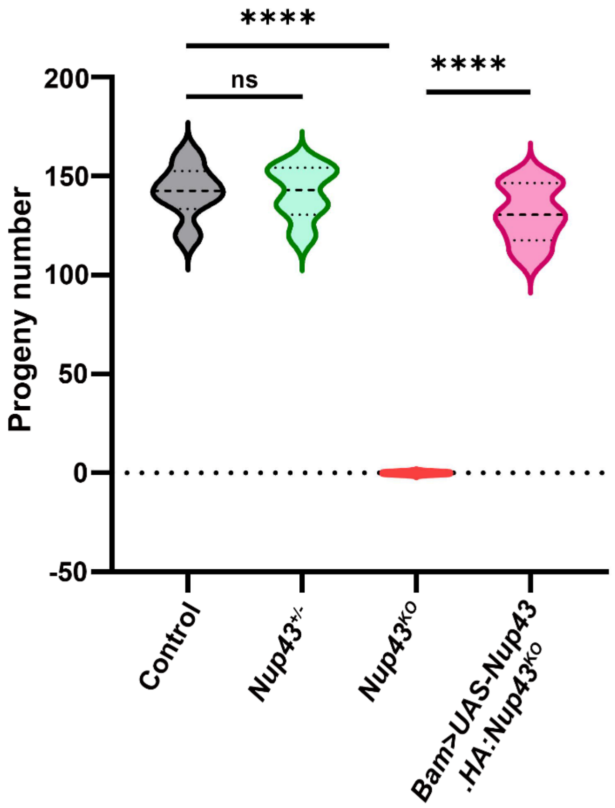

**Fig. S5. Germ-line specific expression of HA-tagged Nup43 transgene rescues male fertility.**

Male fertility assay of control (*w*$^{1118}$), heterozygous Nup43 mutant (*Nup43*$^{+/-}$), homozygous Nup43 mutant (*Nup43*$^{KO}$), and Nup43 transgene rescue (*Bam.Gal4>UAS-Nup43.HA: Nup43*$^{KO}$) males crossed with wild-type (*w*$^{1118}$) females at 25 °C. Subsequently, the number of resulting adult F1 progeny was counted. Data are represented from at least three independent experiments. Statistical significance was derived from the Student's t-test. Error bars represent SEM. Asterisks indicate significance levels. ****p = <0.0001 and ns is non-significant.

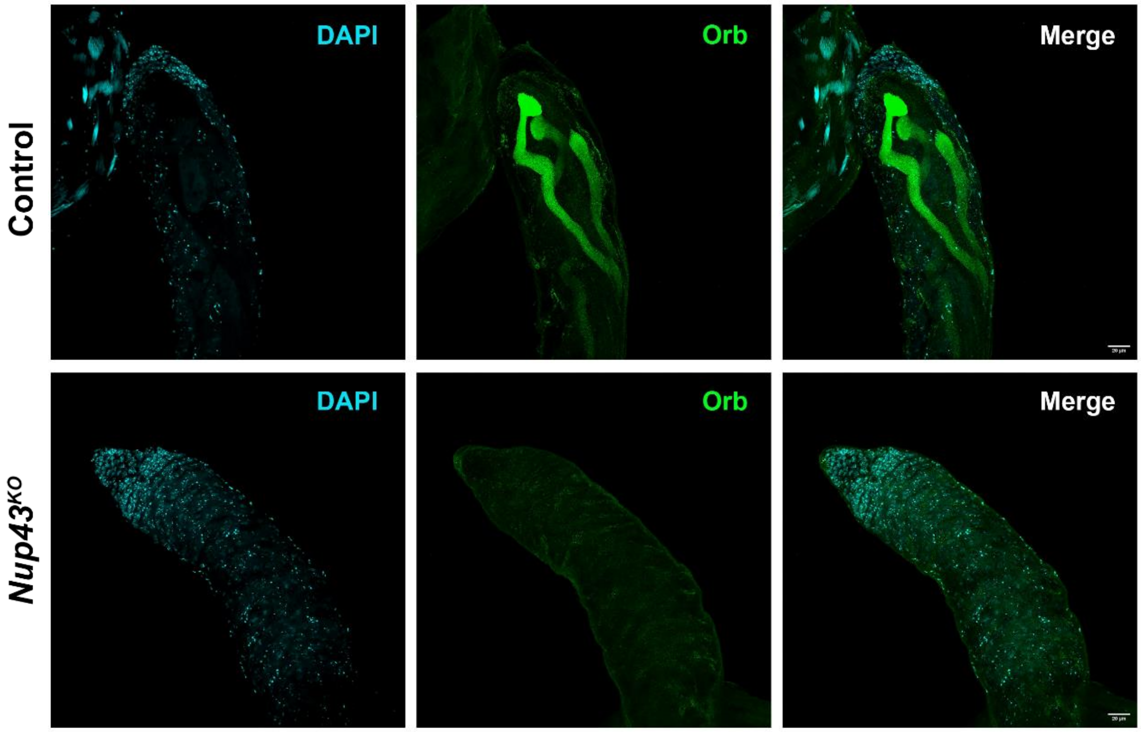

**Fig. S6. The orb staining is missing from the caudal end of the sperm axoneme in the Nup43 mutant testis.**

Immunostaining of the testis of control (*w*$^{1118}$) and *Nup43*$^{KO}$ with orb (green) to visualize sperm axoneme. Chromatin stained with DAPI (cyan). The scale bar represents 20 µm.

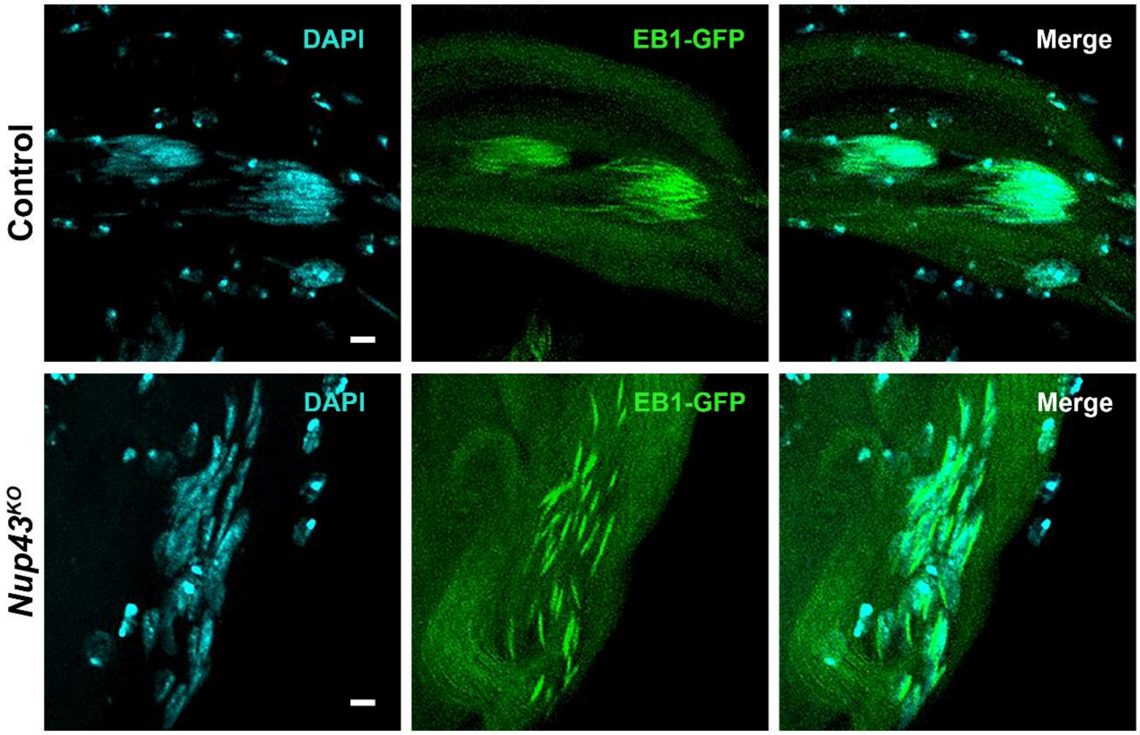

**Fig. S7. The organization of microtubules remains unaffected in Nup43 mutants.** Images of spermatid cysts from control (*EB1-GFP*) testes and the *Nup43*$^{KO}$ (*EB1-GFP; Nup43*$^{KO}$), microtubule binding EB1-GFP marks spermatids. Chromatin stained with DAPI (cyan) to visualize spermatid nuclei. The scale bar represents 5 µm.

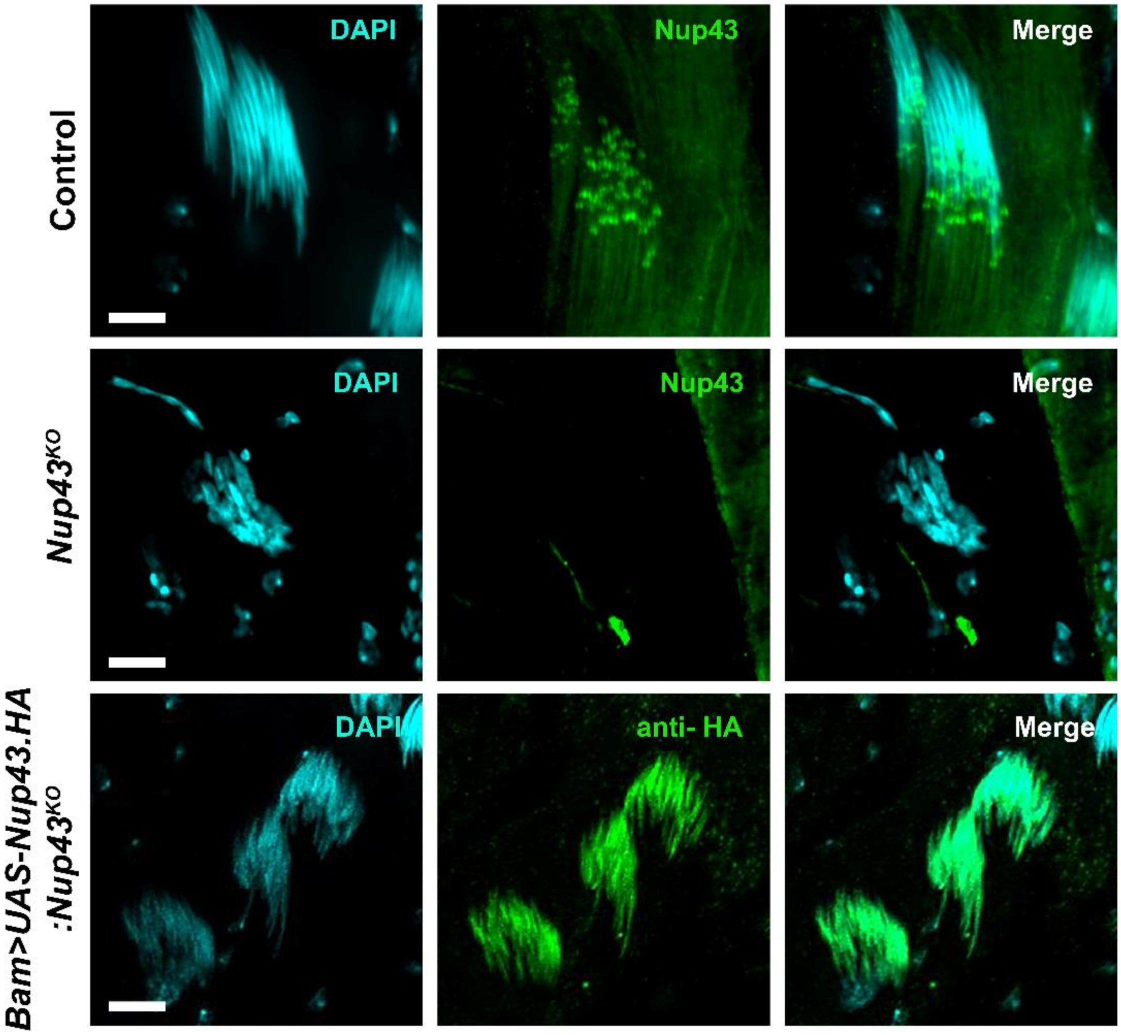

**Fig. S8. Nup43 localizes around individualization complexes.** Immunostaining of the control (*w*[1118]) and *Nup43*[KO] testes (top and middle panels, respectively) with Nup43 staining (green). Immunostaining of the rescue testes (*Bam.Gal4>UAS-Nup43. HA: Nup43*[KO]) with anti-HA (green, bottom panels). Chromatin stained with DAPI (cyan) to visualize spermatid nuclei. The scale bar represents 5 µm.

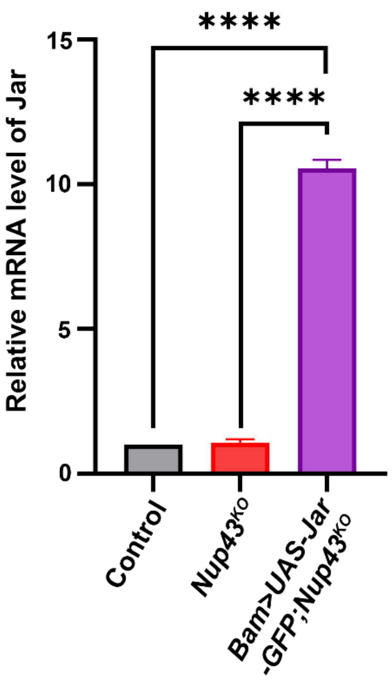

**Fig. S9. *Nup43$^{KO}$* mutants have highly increased levels of jar transcript.** The graph shows quantification of jar transcript levels from control (*w$^{1118}$*), *Nup43$^{KO}$*, and *Bam>UAS-Jar-GFP; Nup43$^{KO}$* testes. Data are represented from at least three independent experiments. Statistical significance was derived from the Student's t-test. Error bars represent SEM. Asterisks indicate significance levels. ****p = <0.0001 and ns is non-significant.

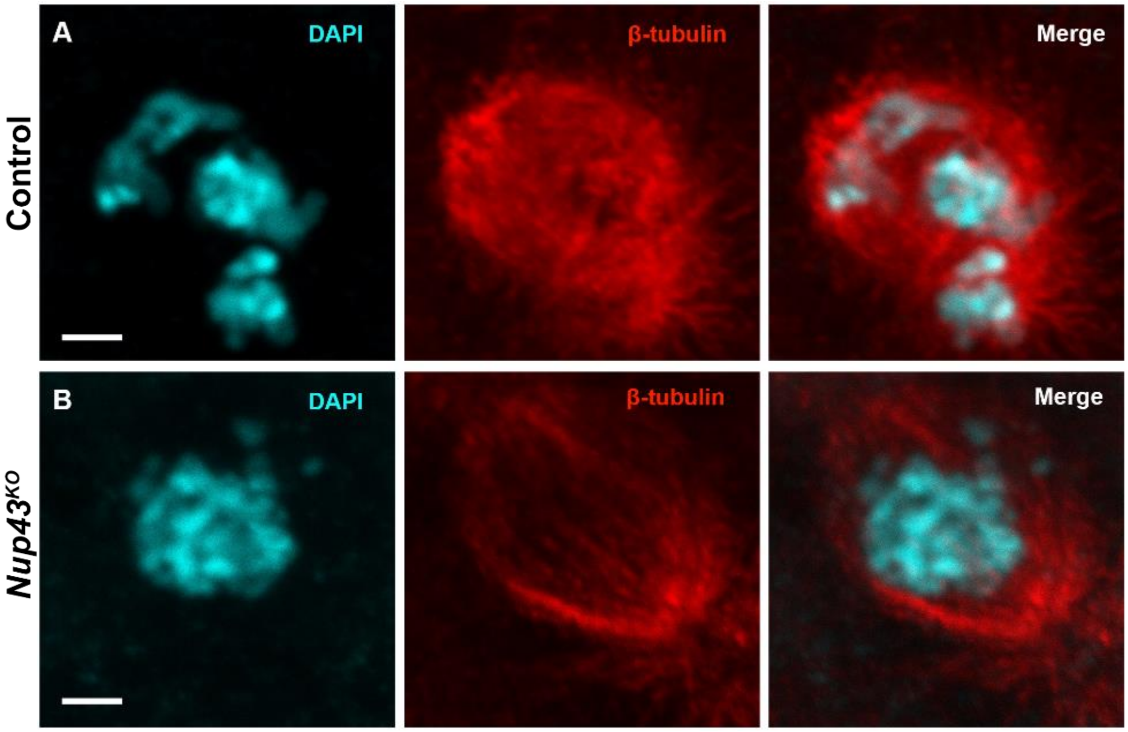

**Fig. S10. Chromosomal alignment as well as mitotic assembly is defective in *Nup43$^{KO}$* mutants during 1st mitotic division.** (A-B) Immunostaining of the embryos after ~ 2 hrs post-fertilization from *Nup43$^{KO}$* mutant (*Nup43$^{KO}$* females mated with *$w^{1118}$* males), showing microtubules labeled with anti-β-tubulin (red). Chromatin is stained with DAPI (cyan) to visualize nuclei. The scale bar represents 2 µm.

**Table S1. Similarity and Identity of Nup43 orthologs to hNup43 protein.**

| **Organism** | **Identity with hNup43 (%)** | **Similarity with hNup43 (%)** | **Query cover (%)** |
|---|---|---|---|
| Mouse | 93.7 | 97.1 | 100 |
| Zebrafish | 61.4 | 75.9 | 99 |
| *Drosophila* | 26.3 | 48.9 | 97 |

**Table S2. Reagents and tools were used in this study.**

| Reagents or Resource | SOURCE | IDENTIFIER |
|---|---|---|
| **Antibodies** | | |
| Rabbit anti-Nup43 | This study | N/A |
| Mouse anti-mAb414 | Biolegend | Cat#902902 |
| Mouse anti- lamin Dm0 | DSHB | Cat# ADL67.10 |
| Mouse anti-orb | DSHB | Cat#6H4 |
| Mouse anti-Ran | BD | Cat#TC4 |
| Mouse anti-β tubulin | DSHB | Cat#E7 |

| | | |
|---|---|---|
| Mouse anti-HA | CST | Cat#2367T |
| Rabbit anti-HA | CST | Cat#3724T |
| Goat Alexa-Fluor Plus 680 | Invitrogen | Cat#A32734 |
| Goat Alexa-Fluor Plus 800 | Invitrogen | Cat#A32730 |
| Goat anti-rabbit Alexa Fluor 488 | Invitrogen | Cat#A11034 |
| Goat anti-rabbit Alexa Fluor 568 | Invitrogen | Cat#A11036 |
| Goat anti-mouse Alexa Fluor 488 | Invitrogen | Cat#A11029 |
| Goat anti-mouse Alexa Fluor 568 | Invitrogen | Cat#A11004 |
| Goat anti-rabbit Alexa Fluor 647 | Jackson ImmunoResearch | Cat#111-605-045 |
| **Chemicals, peptides, and recombinant proteins** | | |
| Fluoroshield | Sigma-Aldrich | Cat#F6057 |
| Phalloidin-647 | Jackson ImmunoResearch | Cat#PHDN1-A |
| TritonX-100 | Sigma-Aldrich | Cat#X100-1L |
| Tween-20 | Sigma-Aldrich | Cat#274348 |
| PBS | In-House preparation | N/A |
| Paraformaldehyde (PFA) | Sigma-Aldrich | Cat#158127 |
| Normal Goat Serum (NGS) | The Jackson Laboratory, USA | Cat#005-000-001 |
| iTaq Universal SYBR® Green Supermix | BIORAD | Cat#1725122 |

| | | |
|---|---|---|
| G9 Taq Polymerase<br><br>10X buffer with $MgCl_2$ | GCC Biotech | Cat#G7115 |
| dNTPs | SBS GENETECH | Cat#EN-2 |
| EDTA | Sigma-Aldrich | Cat#03690 |
| SDS | HIMEDIA | Cat#GRM886 |
| Proteinase-K | MP Biomedicals | Cat#193981 |
| Tris | HIMEDIA | Cat#MB029 |
| Potassium acetate | HIMEDIA | Cat#MB042 |
| Isopropanol | MP Biomedicals | Cat#194006 |
| Ethanol | Merck | Cat#100983 |
| Sucrose | ANJ Biomedicals | Cat#100314 |
| IPTG | Sigma-Aldrich | Cat#I6758 |
| Protease inhibitor Cocktail | Roche | Cat#04693132001 |
| Lysozyme | SRL | Cat#45822 |
| NaCl | Emparta ACS | Cat#1.93206.0521 |
| Sodium deoxycholate | Sigma-Aldrich | Cat# 30970 |
| EGTA | Sigma-Aldrich | Cat#E4378 |
| Sodium azide | Sigma-Aldrich | Cat#438456 |
| Urea | MP Biomedicals | Cat#194857 |
| Sodium Hypochlorite Solution (Liquid Bleach) | SRL | Cat#25366 |
| n-Heptane | Honeywell (Fluka) | Cat# 34495 |
| **Critical commercial assays** | | |

| | | |
|---|---|---|
| RNA isolation kit | Favorgen Biotech | Cat#FATRK-001-2 |
| iScript$^{TM}$ cDNA synthesis kit | BIORAD | Cat#170-8891 |
| **Experimental models: Organisms/strains** | | |
| *Drosophila melanogaster: Nup43 gRNA line* | C-CAMP, Bangalore, India | N/A |
| *Drosophila melanogaster: y[1] M{w[+mC]=nanos-Cas9.P}ZH-2A w[*]* | Bloomington *Drosophila* Resource Center | BL-54591 (nanos.Cas9) |
| *Drosophila melanogaster: w[*]; P{w[+mC]=His2Av-mRFP1}II.2* | | BL-23651 (Histone2Av-mRFP) |
| *Drosophila melanogaster: w[*]; P{w[+mC]=protamineB-eGFP}2/CyO; P{w[+mC]=dj-GFP.S}3/TM3, Sb[1]* | | BL-58406 (ProtamineB-eGFP) |
| *Drosophila melanogaster: w[*]; P{w[+mC]=dj-GFP.S}AS1/CyO* | | BL-5417 (dj-GFP) |
| *Drosophila melanogaster: y[1] w[*] P{w[+mC]=bam-GAL4:VP16}1* | | BL-80579 (Bam-Gal4) |
| *Drosophila melanogaster: M{UAS-Nup43.ORF.3xHA.GW}ZH-86Fb* | Gift from Prof. Udai Bhan Pandey | FlyORF-F003133 (UAS-Nup43.HA) |
| *Drosophila melanogaster: w[*]; P{w[+mC]=Mst27D.mCherry}II.3* | | BL-95385 (Mst27D.mCherry) |

| | | |
|---|---|---|
| *Drosophila melanogaster: w[*]; P{w[+mC]=Eb1-GFP.8322}2* | Bloomington *Drosophila* Resource Center | BL-99926 (Eb1.GFP) |
| *Drosophila melanogaster: w[*]; P{w[+mC]=UAS-jar.GFP}2/SM6a* | | BL-67606 (UAS-jar.GFP) |
| *Drosophila melanogaster: Nup43$^{KO}$/TM6.Tb* | This study | N/A |
| *Drosophila melanogaster: dj-GFP/Cyo; Nup43$^{KO}$/TM6.Tb* | | |
| *Drosophila melanogaster: dj-GFP/Cyo; UAS-Nup43.HA:Nup43$^{KO}$/TM6.Tb* | | |
| *Drosophila melanogaster: Bam-Gal4;+; Nup43$^{KO}$/TM6.Tb* | | |
| *Drosophila melanogaster: Eb1-GFP/Cyo; Nup43$^{KO}$/TM6.Tb* | | |
| *Drosophila melanogaster: Mst27D.mCherry/Cyo; Nup43$^{KO}$/TM6.Tb* | | |
| *Drosophila melanogaster: UAS-jar.GFP/Cyo; Nup43$^{KO}$/TM6.Tb* | | |
| *Drosophila melanogaster: UAS-Nup43.HA:Nup43$^{KO}$/TM6.Tb* | | |
| *Drosophila melanogaster: ProtamineB-eGFP:His2Av-mRFP/Cyo* | | |
| *Drosophila melanogaster: ProtamineB-eGFP:His2Av-mRFP/Cyo; Nup43$^{KO}$/TM6.Tb* | | |

| Oligonucleotides | | |
|---|---|---|
| Primer: Nup43_gRNA1: 5'-<br><br>CACGCACTACATATCCGAGAAGG-3' | This Study | N/A |
| Primer: Nup43_ gRNA2: 5'-<br><br>CATGGAAGTCGTCTGGTCTGCGG-3' | | |
| Primer: 5'UTR Primer1_F: 5'-<br><br>GCTATTGAAAAATTGGCGCC-3' | | |
| Primer: 3'UTR Primer1_R: 5'-<br><br>GCCTAATATGCATTGAATCC-3' | | |
| Primer: 5'UTR Primer2_F: 5'-<br><br>CCGCTAAGATATGTTGCGAT-3' | | |
| Primer: 3'UTR Primer2_R: 5'-<br><br>CCCATAAACACCTGTATACG-3' | | |
| Primer: Nup43_RT_F: 5'-<br><br>CAGCCGAAGACCACCTTTAT-3' | | |
| Primer: Nup43_RT_R: 5'-<br><br>CCTATCTCGTTGATGGGACTTT-3' | | |
| Primer: jar_RT_F: 5'-<br><br>GGTGTACCACGCCTGGAAGG-3' | | |
| Primer: jar_RT_R: 5'-<br><br>GCTGGGCGGTCACAATCTCC-3' | | |
| Primer: Rpl49_RT_F: 5'-<br><br>CGTTTACTGCGGCGAGAT-3' | | |

| | |
|---|---|
| Primer: Rpl49_RT_R: 5'-GTGTATTCCGACCACGTTACA-3' | |
| **Software and algorithms** | |
| ImageJ/FIJI | National Institute of Health, USA |
| GraphPad Prism Software | GraphPad |
| Adobe Photoshop 2023 | Adobe |
| QuantStudio Design & Analysis Software | Applied Biosystems |
